# Multi-Modal Mass Spectrometry Imaging Reveals Isomer-Specific PFOS Distribution and Spatial Lipid Disruption in Mouse Liver

**DOI:** 10.1101/2025.10.13.682167

**Authors:** Aidan J. Reynolds, Jaide A. Mickel, Srijana Shrestha, Rance Nault, Tian (Autumn) Qiu

## Abstract

The linear and branched isomers of perfluorooctane sulfonate (PFOS), a legacy per-and polyfluoroalkyl substances (PFAS), show differential bioaccumulation and toxicological responses. However, their spatial distribution within tissues and their effects in spatial contexts remain insufficiently resolved. In this work, we present a multi-modal mass spectrometry imaging (MSI) strategy to resolve the on-tissue distribution of linear *(n-*) and branched (*br-*) PFOS isomers in mouse liver and PFOS-induced spatial lipid disruption. Liver samples from exposed mice were analyzed by multi-modal MSI, integrating matrix-assisted laser desorption/ionization-trapped ion mobility spectrometry (MALDI-TIMS)-MSI and post-MALDI immunofluorescence of canonical zonation markers. We found that PFOS exhibited dose-dependent and zonation-specific distribution, with enrichment in pericentral regions compared to periportal. *br-*PFOS displayed a more homogeneous distribution compared to *n-*PFOS. Spatial lipidomics revealed concomitant zonal disruptions in hepatic lipid abundance and distribution, reflecting zonal lipid disruptions not accessible by bulk measurements. Together, this study establishes a multi-modal, spatial-resolved analytical workflow for mapping on-tissue PFOS isomer distribution and lipid disruptions. By demonstrating that PFOS isomeric structure governs tissue localization and is associated with zonal lipid disruptions, our findings provide new insights into the spatial dimension of toxicokinetics/ toxicodynamics and underscore the need for isomer-resolved, spatial-focusing approaches on environmental and toxicology research.

**TOC Figure:** 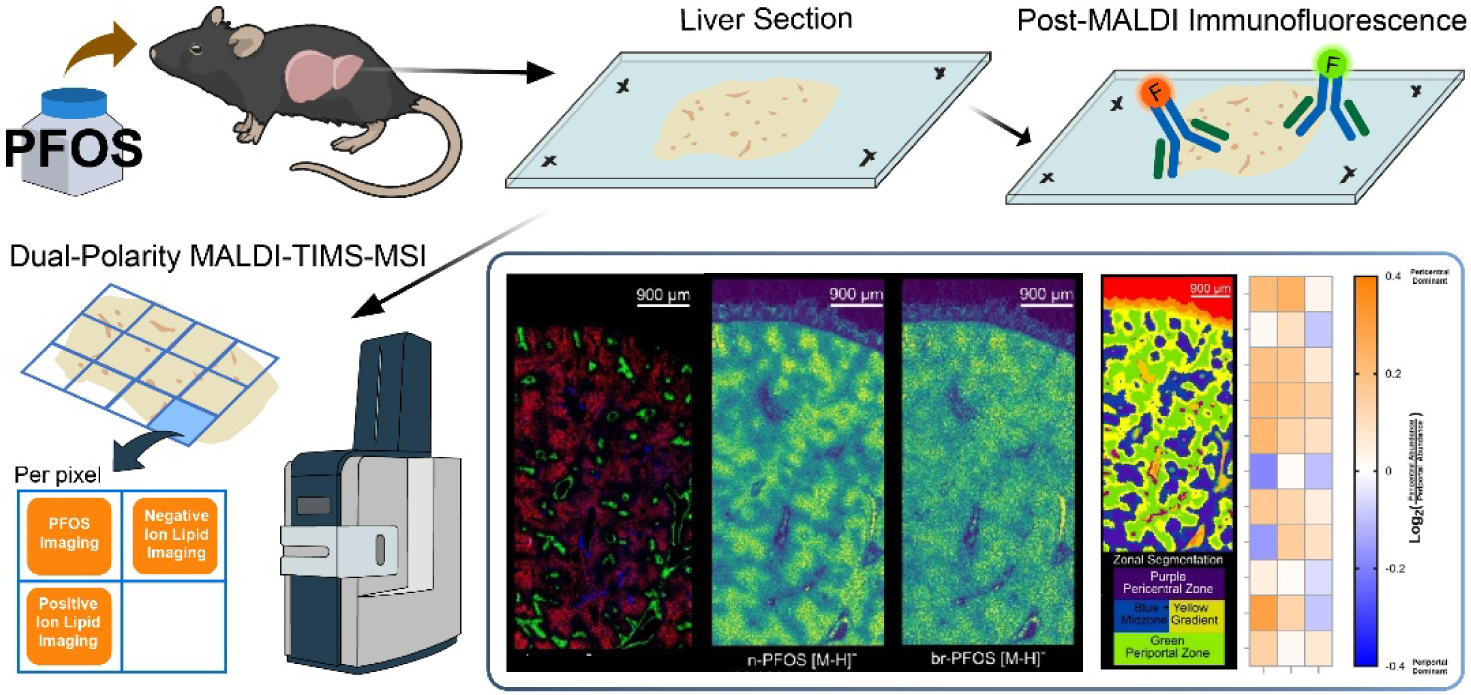

*Synopsis:* Multimodal mass spectrometry imaging reveals isomer-specific PFOS distribution and zonal lipid disruption in liver, linking chemical structure to spatial toxicology of PFAS.

## Introduction

Per-and polyfluoroalkyl substances (PFAS) are emerging environmental contaminants that contain at least one fully fluorinated methyl or methylene carbon atom.^1^ Their carbon-fluorine bonds provide exceptional stability, enabling widespread industrial and commercial use while also earning them the label "forever chemicals" due to their resistance to degradation and persistence in the environment and human bodies.^2,3^ The distribution and accumulation of PFAS in tissues is a critical component of their biological fate *in vivo*, as it dictates the extent of direct interaction of PFAS with organs and contributes to toxic responses.

Strong evidence by previous studies have proven the links between PFAS compound structures and PFAS exposure outcomes, including distribution and accumulation in tissues.^3–5^ One largely under-exploited structural feature is the isomeric structures of PFAS, namely linear vs. branched structures. With the advance of analytical methods, evidence has shown a wide existence of branched PFAS isomers from sources to applications.^6^ For example, the widespread perfluorooctane sulfonate (PFOS) products usually contain about 30% of the mixed branched PFOS isomers, a non-negligible fraction, from electrochemical fluorination production.^7^ Notably, the linear and branched isomeric PFAS structures have shown differential cell uptake and are associated with differential toxicity outcomes and health effects.^6,8,9^ Thus, neglecting isomeric PFAS structures can significantly compromise our capability to accurately assess the toxicity and risk of PFAS for best regulatory practices.

Previous reports showed isomer-specific binding affinities of PFOS to serum proteins and potentially different fractionations between blood plasma and cells.^10–12^ A recent report showed that total branched PFOS preferably accumulated or concentrated over linear PFOS in rat tissues.^13^ However, previous research on isomeric PFAS biodistribution failed to address one important factor: the spatial heterogeneity in biological tissues. Biological tissues are highly heterogeneous across scales. Xenobiotic accumulation in tissues is highly influenced by the tissue type, molecular composition, as well as the physio-chemical properties of contaminants.^14,15^ Mass spectrometry imaging (MSI) technology enables spatially resolved, on-tissue mapping of analytes for spatial-omics studies. Using various MSI technologies, previous work has revealed heterogeneous distribution of PFOS and other PFAS compounds in animal tissues,^16–21^ yet the isomeric structures were not resolved on tissue in those studies.

PFOS exposure has been linked to perturbed lipid homeostasis and hepatic steatosis.^22,23^ Focusing on liver, a primary detoxification organ, previous studies have revealed disruption in hepatic lipid metabolism from PFOS exposure via MS-based lipidomics,^18,24^ gene expression and molecular assays,^23^ and proteomics.^25^ Using liquid chromatography (LC)-MS lipidomics, Li et al. reported altered levels upon PFOS exposure for a wide panel of hepatic lipids, including lysophosphatidylcholine (LPC), phosphatidylcholine (PC), ceramides (Cer), phosphoinositols (PI), triacylglycerol (TG), sphingomyelin (SM) and more.^18^ Notably, hepatic lipids show spatial distribution patterns mapped to liver zonation, a honeycomb-like pattern of hepatic functional units (lobules). Each lobule is bounded by portal triads consisting of the portal vein, hepatic artery, and bile duct that drain into a central vein at the middle creating regions with distinct microenvironments and metabolic function.^26–28^ As a previous study showed that PFOS can promote cellular changes and lipid accumulation in centrilobular hepatocytes,^29^ the importance of characterizing hepatic zonation-specific alterations in lipid metabolism needs to be addressed.

Herein, we for the first time report the observation of isomer-specific distribution of linear (n-) vs. summed branched (br-) PFOS in liver of exposed mice and characterized the zonation-specific alterations in hepatic lipid abundance using matrix-assisted laser desorption/ionization-trapped ion mobility spectrometry-mass spectrometry imaging (MALDI-TIMS-MSI). An illustration of experimental and analysis pipeline is illustrated in **Figure 1**. Our previous report demonstrated detection of PFOS using MALDI-TIMS-MS analysis.^30^ In this study, mice were exposed to commercially sourced PFOS via oral administration, followed by liver tissue collection, embedding, freezing, and cryo-sectioning for multi-modal imaging workflow that integrated autofluorescence, dual polarity MALDI-TIMS-MS imaging of PFOS and lipids, and post-MALDI immunofluorescence staining on the same tissue. We successfully separated linear and branched PFOS isomers on-tissue using TIMS and discovered that while both n- and br-PFOS shows pericentral distribution, br-PFOS was relatively more homogenously distributed compared to n-PFOS. With laser offset, dual polarity MSI was performed on the same tissue to acquire data on endogenous lipids. Results showed that PFOS exposure induced changes in hepatic lipid dysregulation, characterized by zonal lipid abundance changes and alterations in lipid zonal distribution. Overall, our results demonstrated the importance of considering linear vs. isomer for PFAS distribution in tissue and the resulted toxicity responses in their spatial contexts.

**Figure 1.**
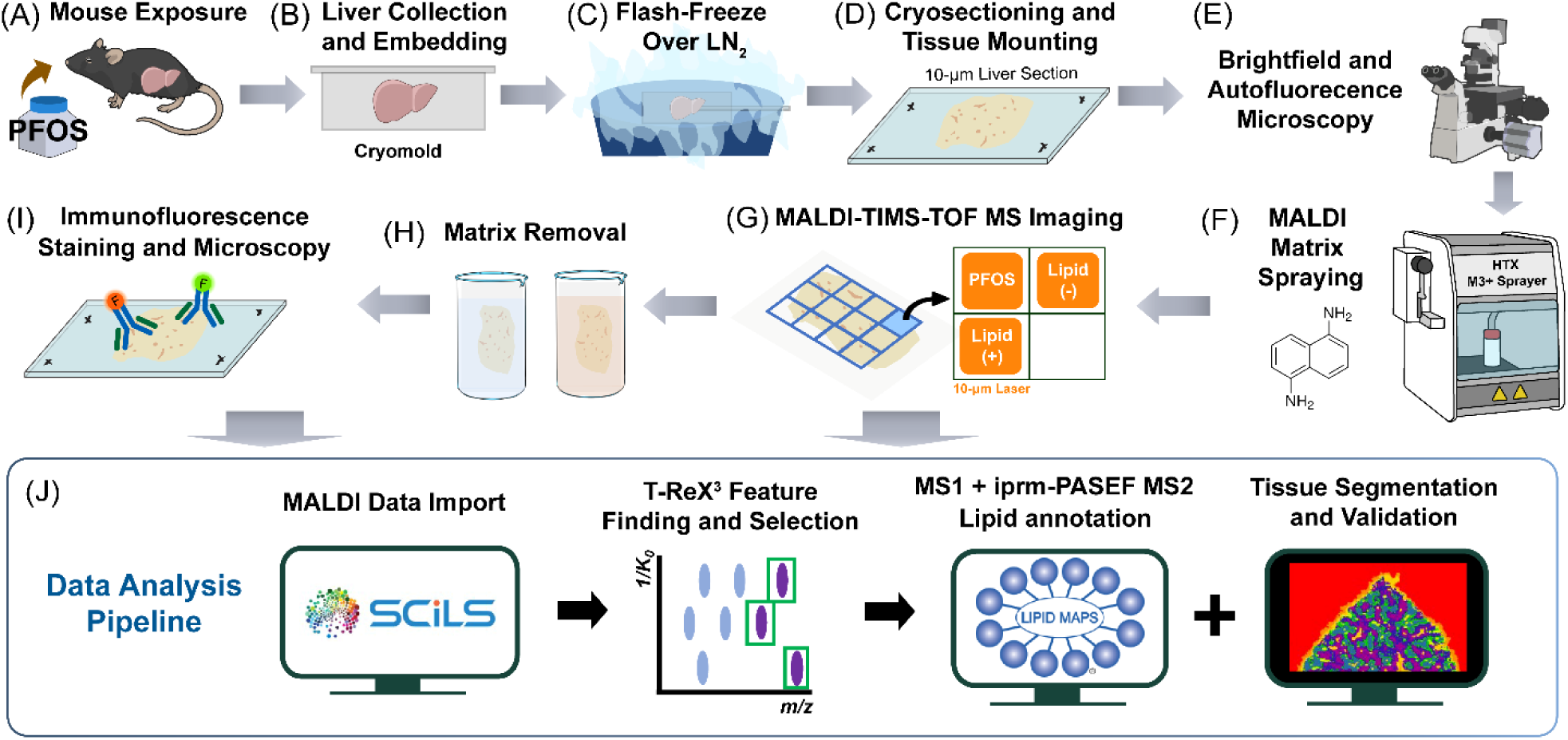
Illustration of experimental and data analysis pipeline for multi-modal MS imaging of PFOS-exposed mouse liver. (A) Mouse exposure with PFOS administered via drinking water; (B) Collection and embedding of liver samples after 3-month exposure; (C) flash-freezing over liquid nitrogen. (D) Cryosectioning of 10 µm sections with 1 sample from each dose group on each slide, (E) detection of tissue area by autofluorescence microscopy; (F) application of MALDI matrix; (G) spatial analysis of PFOS isomers and lipid species by MALDI-TIMS-TOF mass spec imaging; (H) removal of MALDI matrix by consecutive ethanol washes; and (I) validation of zonation by post-MALDI immunofluorescence imaging of known markers; (J) data analysis pipeline, including feature finding, metabolite annotation, and segmentation of lobular zonation.

## Results

### Detection and characterization of linear and branched PFOS on liver tissue

Under negative mode analysis, PFOS on liver tissue was detected primarily as deprotonated ions, [M-H]^-^, consistent with our previous report on pure PFOS standards (**Figure 2A-C**).^30^ We previously reported nearly baseline separation for three PFOS structural isomers with 500 and 800 ms as TIMS ramp time.^30^ However, longer ramp time significantly increases the imaging acquisition time. Thus, we optimized the ramp time for TIMS and decided on 150 ms to balance the needs for both isomer separation and acquisition time. Two PFOS isomer peaks were successfully identified on tissue, with collision cross section (CCS) determined as 166.9 ± 0.8 and 163.0 ± 0.8 at *m/z* 498.9313 ± 0.0003 (average ± standard deviation of four replicates of exposed samples). Previous report using drift tube IMS coupled with LC-MS determined the CCS for linear PFOS to be 167.9, while branched PFOS structures range from 161.3 to 166.1 for different branched structures.^31^ From testing analytical standards of several PFOS isomers and matching their spectra, we determined that the major PFOS isomer peak at 166.9 ± 0.8 to be n-PFOS, and the minor PFOS isomer peak at 163.0 ± 0.8 to be a mixture of branched PFOS that consist of at least P3MHpS and P4MHpS, based on observed diagnostic fragment ions and measured CCS values from MALDI-TIMS-TOF MS/MS via iprm-PASEF mode analysis **(Tables S1 and S2, Figures S1 and S2)**.^32^ More detailed discussions regarding characterization can be found in Supplemental Information.

**Figure 2.**
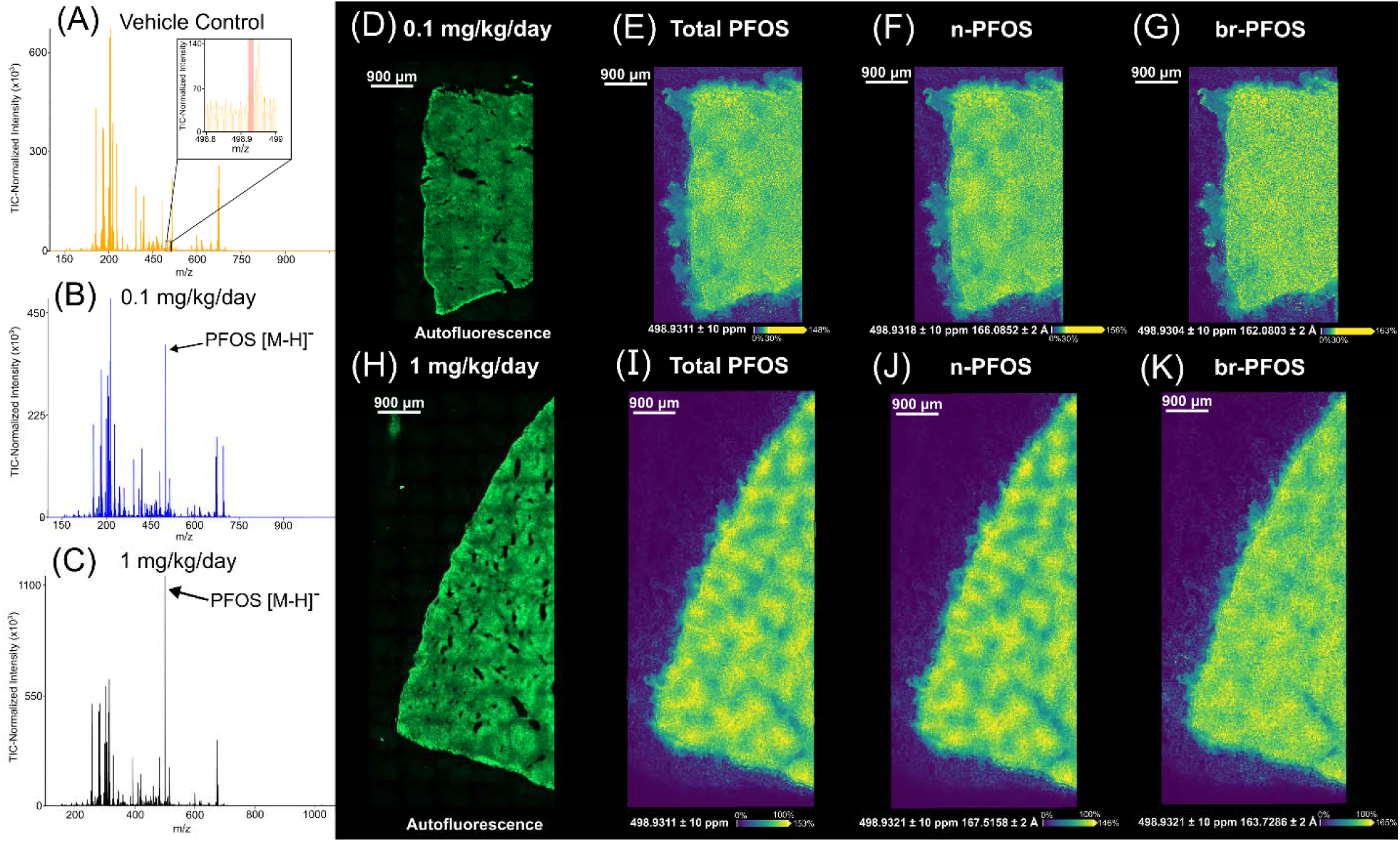
Detection and spatial distribution of PFOS on exposed liver tissue sections. (A-C) average spectra of vehicle control and PFOS-exposed liver tissue sections. The signals of PFOS ([M-H]^-^) are indicated by arrows on the spectra. (D-G) Pre-MALDI brightfield and green autofluorescence providing tissue area overview and MALDI-MSI ion images for total, linear, and branched PFOS from 0.1 mg/kg/day PFOS-exposed liver samples. (H-K) Images from 1 mg/kg/day PFOS-exposed liver samples. Scale bars in ion images show relative scales; therefore, pixel colors should not be directly compared among ion images. MALDI-MSI parameters: matrix-DAN; polarity-negative; lateral spatial resolution-20 μm; other parameters-see Experimental methods.

### Dose-dependent and isomer-specific localization of PFOS to hepatic zonation

Average mass spectra of liver regions showed that PFOS accumulation in liver was dose-dependent, as the PFOS peak in 1 mg/kg/day PFOS-exposed mouse livers was significantly higher than the PFOS peak in 0.1 mg/kg/day dosed animals (**Figure 2A-C**). Such trend was also shown in Figure S3 where liver PFOS images from two doses are plotted on the same intensity scale (**Figure S3**). To better show the distribution patterns in low-dose tissues, we adjusted the intensity threshold for the scale bar as shown in **Figure 2E-G**. Autofluorescence images (**Figure 2D, H; Figure 3A**) outline tissue morphology. Ion images of total PFOS (**Figure 2E, I**) and linear PFOS (**Figure 2F, J**) showed distribution patterns that resemble liver zonation in both 0.1 and 1 mg/kg/day exposed samples. Interestingly, in the 0.1 mg/kg/day exposed livers, branched PFOS demonstrated a more homogenous distribution without clear specific patterns, while a similar zonation-like pattern showed up, although not as significant as linear PFOS, in 1 mg/kg/day exposed livers (**Figure 2G, K**). We confirmed that no PFOS was detectable in imaged vehicle control tissues (**Figure S4**), and such PFOS distribution patterns are consistent amongst all replicates of PFOS-exposed liver samples (**Figure S5**).

**Figure 3.**
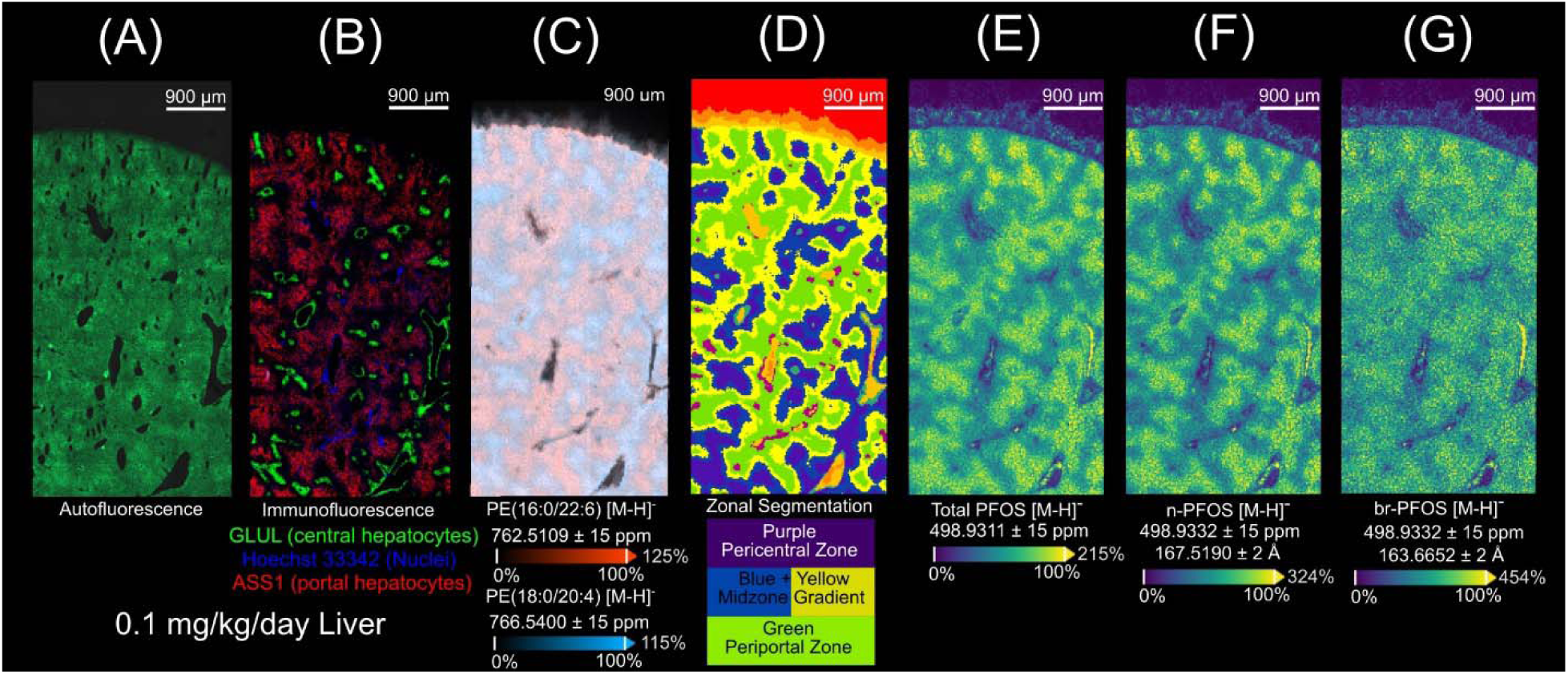
Multi-modal imaging results on the same 0.1 mg/kg/day PFOS-exposed mouse liver tissue section. (A) Pre-MALDI brightfield and GFP autofluorescence used as tissue area overview. (B) Post-MALDI immunofluorescence imaging of zonal hepatocyte markers used for validation of zonal segmentation. Livers were stained for Hoechst 33342 (Nuclei; blue), ASS1 (portal hepatocytes; red), and GLUL (central hepatocytes; green). (C) Periportal and pericentral distribution of PE(16:0/22:6) and PE(18:0/20:4), respectively, putatively annotated via accurate mass matching and confirmed with on-tissue MS/MS. (D) Zonal segmentation based on lipidomic profiles of imaged tissue regions visualizing pericentral zone (purple), midzone gradient (blue and yellow), and periportal zone (green). Additional colors represent non-tissue background, tissue edges, and vessels, and were not used for analysis in this study. (E-G) MALDI-MSI ion images for total, linear and branched PFOS. A representative section from the 1 mg/kg/day dose group is shown in **Figure S4**. MALDI-MSI parameters: matrix-DAN; polarity-negative; lateral spatial resolution-20 μm; other parameters-see Experimental.

To confirm whether PFOS distribution is related to liver zonation, we performed a multi-modal workflow on the same liver tissue section to overlay MS imaging data with post-MALDI immunofluorescence staining for zonal hepatocyte markers. Examples of immunofluorescence images are shown in **Figure 3B** and more images can be found in **Figure S5-S7**. Liver sections were stained for Hoechst 33342 (Nuclei; blue), ASS1 (portal hepatocytes; red), and GLUL (central hepatocytes; green). By overlaying ion images with immunofluorescence images for the same region, it is evident that the total and linear PFOS localize in the pericentral area (**Figure 3E-F** for 0.1 mg/kg/day, **Figure S7E-F** for 1 mg/kg/day). Similar to **Figure 2G and 2K**, the branched PFOS showed a more homogeneous pattern (**Figure 3G**). We also demonstrate periportal and pericentral distribution of two endogenous lipids, annotated by accurate mass and on-tissue MS/MS as PE(16:0/22:6) and PE(18:0/20:4) (**Figure 3C**). The pericentral PE(18:0/20:4) co-localize well with PFOS signals, further confirming the pericentral distribution of total and linear PFOS.

To enable zonation-specific analysis of PFOS, we performed unsupervised segmentation using bisect *k*-means clustering to cluster pixels based on the features identified from MS1 spectra; an example of the segmentation process is shown in **Figure S8**. Results from segmentation were validated against immunofluorescence of canonical zonation markers (**Figure 3B, D; Figure S9**). While mid-zone regions were also identified, we decided to focus on pericentral and periportal regions to reflect the most disparate regions of the lobule. Two samples were excluded from quantitative analyses as livers were clearly outliers in their histopathological features which may stem from a ∼10% incidence of congenital portal shunts in C57BL/6J mice (**Figure S10**).^33^ To ensure that all PFOS intensities used for downstream statistical analyses was quantifiable, signal-to-noise (S/N) ratios were taken from low-intensity areas within segmented pericentral and/or periportal areas of imaged 0.1 mg/kg/day liver samples. Across all replicates from randomly sampled regions with low-intensity PFOS signal, both n-PFOS and br-PFOS signals were found to have S/N > 10, confirming that entire segmented areas could be used for quantitative analyses (**Figure S11**). On-tissue ionization efficiency experiments were also conducted to confirm if branched isomers yielded similar signal intensity to n-PFOS. We found that at equivalent concentrations of 500 ng/mL, n-PFOS, P3MHpS, P4MHpS, P5MHpS, and P6MHpS yielded similar signal intensities suggesting that they have equivalent ionization efficiencies (**Figure S12**).

Intensities of PFOS isomers were extracted in segmented regions representing pericentral and periportal zones, and a pericentral:periportal (PC:PP) ratio was calculated to represent zonal distribution (**Equation 1**). A log2-transformed PC:PP ratio was calculated so that zero means no zonal preference, positive value means pericentral preference, and negative value means periportal preference. Results showed that both n- and br-PFOS exhibited a preference to accumulate in the pericentral zone at both 0.1 and 1 mg/kg/day exposure level, while br-PFOS showed less zonal preference towards a more homogenous distribution, characterized by lower PC:PP ratios (**Figure 4**). In addition to cross-animal comparisons, we also performed within-sample comparisons of PFOS distribution between the pericentral and periportal regions within each PFOS-exposed tissue using a conditional autoregressive (CAR) model to correct pixel-wise spatial dependence (**Figure S13, Figure S14**), and results showed that the difference in lobular distribution was evident within each individual samples.^34^ Therefore, results from both within-tissue and cross-animal comparisons together showed that linear and branched PFOS display different spatial distribution patterns in terms of liver zonation, matching our visual evaluation of ion images (**Figure 2 and 3, Figure S5**).

**Figure 4.**
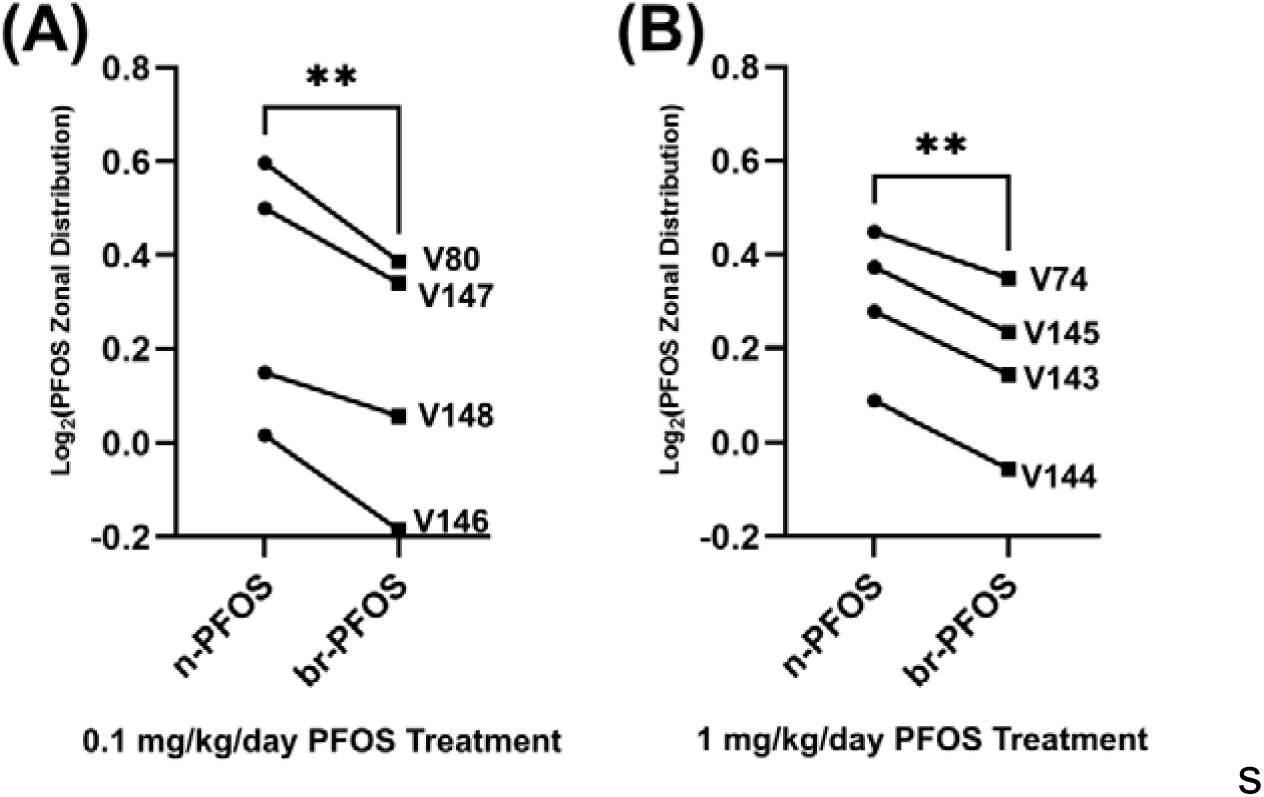
Log_2_ transformed pericentral:periportal ratio analysis of n-PFOS and br-PFOS isomers between pericentral and periportal zones. Zero means no zonal preference, positive value means pericentral preference, and negative value means periportal preference. (A) PFOS isomer zonal distribution within 0.1 mg/kg/day PFOS treatment livers. (B) PFOS isomer zonal distribution within 1 mg/kg/day PFOS treatment livers. Animal IDs are indicated to the right of the paired measurements. Paired Student’s t-test was used as values of each n- and br-PFOS data obtained on the same tissue section. **=0.01.

### PFOS exposure-induced changes in lipid abundance and zonal distribution

PFOS exposure has been reported to disrupt lipid metabolism in the liver.^24,35^ Using 10-μm laser spot and 20-μm raster size, in addition to PFOS analysis, we were able to use two out of four quadrants to map liver metabolite and lipid distribution at both positive and negative polarities (**Figure 1**). Average spectra for lipid imaging under both negative and positive modes from control liver samples are shown in **Figure S15**. From negative mode, a total of 305 tissue-localizing features were found, of which 162 unique lipids were annotated with different levels of confidence. From positive mode, a total of 144 tissue-localizing features were found, of which 112 had matches and 86 lipids were uniquely annotated after deisotoping and collapsing adducts. A complete list of discovered features and their annotations can be found in Supplemental Materials.

For positive mode, we found zonated patterns of lipid species such as PC 34:1 and PC 37:3 in periportal and pericentral zones (**Figure S16**), similarly reported in literature.^27^ When generating segmentation maps using lipid features in positive mode, we found that the feature lists struggled to generate adequate segmentation maps that accurately reflected the zonated pattern of the liver in the high-dose (1 mg/kg/day) groups. Combined with the observed signal variations amongst replicates (**Figure S15B**), in this work, we decided to focus on the results from negative mode analysis which were more consistent across replicates and generated more features. For negative mode, a total of 80 lipids were selected based on intensities and putative MS1 matching for on-tissue MS/MS with iprm-PASEF, out of which 56 lipids were deemed suitable for statistical analysis based on intensity thresholds, image quality, and reproducibility across replicates. MS/MS via iprm-PASEF yielded on-tissue MS/MS data for 41 lipid species (**Table S3**). The annotation was also supported by previous reports on liver lipidomics.^27,28^

We first evaluated zonal lipid abundance changes upon PFOS exposure, aiming to explore zonation-specific lipid biomarkers for PFOS exposure. With zone regions defined by unsupervised segmentation and validated by immunofluorescence, we obtained region-specific, TIC-normalized intensities of endogenous lipids and calculated fold changes (FC) between treated and control groups within each region (**Equation 2**), and a log2-transformation was implemented to normalize the FC before statistical analysis. Heatmaps of log2(FC) revealed PFOS-induced lipid abundance changes in both pericentral and periportal zones for all 56 lipids from negative mode analysis (**Figure S17**). By comparing zona lipid abundances, we found that the trend for up-or down-regulation generally does not differ between low (0.1 mg/kg/day) and high (1 mg/kg/day) dose, where some ions showed dose-dependent FC trends, such as PS(18:0/20:4), and some ions showed similar FC trends in low and high doses like PI(16:0/18:2). Interestingly, some lipids showed opposite trends for low and high doses, such as PI(18:1/20:4). While lipid abundance changed in both zones, changes in abundance showed similar trends of varying magnitudes in both the pericentral and periportal regions (**Figure S17**).

We noticed that despite similar trends of lipid abundance change in both zones, the magnitude of changes seemed to subtly differ. Therefore, we implemented subsequent zonal ratio analysis which revealed shifts in lipid zonal distribution upon PFOS exposure (**Equation 1, Figure 5**). Similarly to defining zonal distribution of PFOS, lipid zonal distribution was quantified by calculating log2-transformed PC:PP ratio of intensity, where zero indicates a lipid has no zonal distribution preference, positive values represent pericentral preference, and negative values represents periportal preference. A full heatmap of all replicates and treatments is presented in **Figure S18**. A hierarchical clustering of both sample and lipid dimensions was performed to seek patterns in lipid ratios. Results showed that all high-dose (1 mg/kg/day) replicates were grouped together, while control and low-dose groups (0.1 mg/kg/day), on the contrary, were not effectively separated (**Figure 5A**). Further clustering effort on only control and low-dose treated groups showed a limited level of separation (**Figure S19**), altogether indicating that with 0.1 mg/kg/day of exposure, the zonal distribution patterns of the 56 lipids we examined are generally similar to control. On the other end, 1 mg/kg/day treatment induced zonal distribution pattern changes in liver. Dunnett multiple comparison for treatment groups and Benjamini-Hochberg FDR for lipid revealed two lipids with statistically significant differences in zonal ratio upon high-dose exposure, and their corresponding ion images of all replicates are shown in **Figure 5B-D**. Both lipids showed pericentral distribution in control and low-dose groups. Upon high-dose exposure, 766.541 (PE 18:0/20:4) moved towards non-preferential distribution, while 814.539 (PE 20:2/22:6) flipped their pattern to periportal.

**Figure 5.**
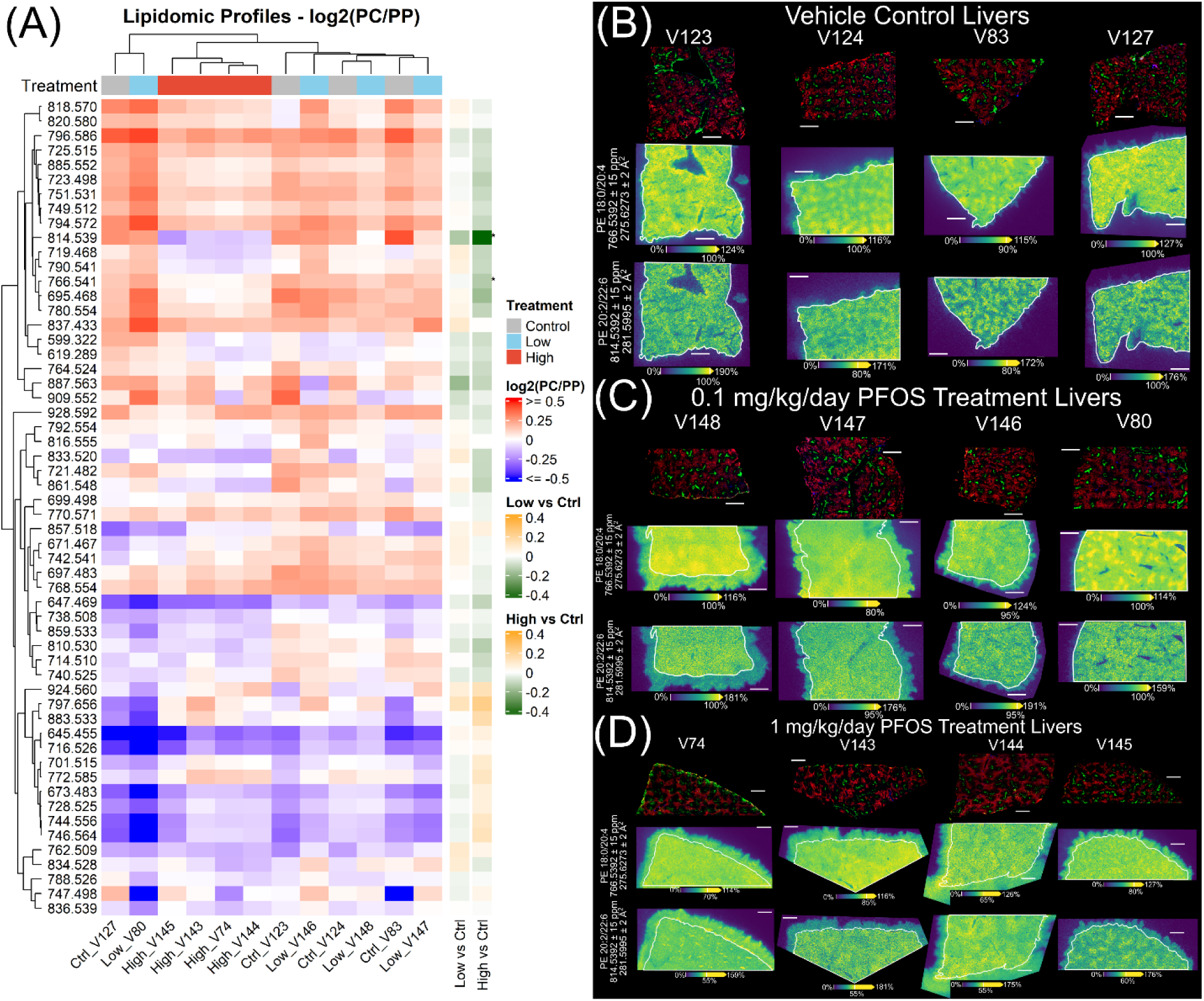
PFOS exposure induced alterations in liver lipid distribution. (A) Hierarchical clustering of groups and lipids. For each lipid feature, zonal distribution was represented as the log2-transformed ratio of PC:PP intensities. Positive values (red scale) indicate pericentral preference. Negative values (blue scale) indicate periportal preference. Zero indicates no preference. Treatment groups are indicated by different colors. For each lipid, effects of treatment on log2(PC:PP) ratios were evaluated using Dunnett’s multiple comparison to compare the low- and high-dose groups separately to the control and Benjamini-Hochberg FDR correction for lipids. Fold changes (low vs ctrl, high vs ctrl) were shown as color bars near the clustering map, and statistical significance was defined as FDR-adjusted q < 0.05 and shown near the fold change bars. (B-D) Ion images of 766.541 (PE 18:0/20:4) and 814.539 (PE 20:2/22:6) in all replicates, together with post-MALDI immunofluorescence imaging of zonal hepatocyte markers. Livers were stained for Hoechst 33342 (Nuclei; blue), ASS1 (portal hepatocytes; red), and GLUL (central hepatocytes; green). The intensity scale bars in ion images were adjusted to improve the visualization of zonal distribution, so the color should not be compared across tissues. White outline in ion images of 1 mg/kg/day PFOS-exposed mouse liver indicates tissue edge. Lipids are annotated via accurate mass matching and on-tissue MS/MS. MALDI-MSI parameters: matrix-DAN; polarity-negative; lateral spatial resolution-20 μm; other parameters-see Experimental methods.

To showcase different kind of pattern changes, **Figure S20** shows AF, IF, segmentation map, and 11 lipid images for all replicates. We noticed that as dose increased, there were progressively pronounced delocalization of lipids beyond tissue edges. Interestingly, PFOS signals (**Figure S5**) did not produce similar levels of delocalization. During sample preparation, we prepared slides where every slide had one control, one low-dose, and one high-dose tissues; we did not observe delocalization effects that were specifically associated with specific slides. Therefore, we believe that the general lipid delocalization pattern we observed is related to exposure-related biological nature of the tissue, for example, related to the elevated lipid droplet content (steatosis) in high-dosed tissues that may cause edge effects during embedding (**Figure S8**).^23,36^ Meanwhile, technical aspects in sample preparation can also contribute to delocalization, particularly when interacting with the biological factors.

## Discussion

By integrating MALDI-TIMS-MS imaging and immunofluorescence, the presented multimodal imaging workflow to our knowledge is the first to enable the separation of structural PFAS isomers on-tissue with direct mapping to hepatic zonation markers. MS imaging techniques have been applied to characterize the distribution of xenobiotics and their metabolites across tissues to better inform pharmacokinetic modeling driven by spatial heterogeneity.^37–40^ However, as PFOS isomers are not readily metabolized, they accumulate in the liver and whether they exhibit different spatial distribution remained unclear. By performing MALDI-MS imaging with a high lateral resolution, we clearly showed spatially heterogenous PFOS distribution in liver sections. Co-localization of PFOS signals with zonation markers from post-MALDI immunofluorescence on the same tissue provided unambiguous evidence that the distribution pattern is zonated and pericentral. We observed similar distribution patterns across four biological replicates.

While the mechanisms for zonal PFOS accumulation and distribution preferences of isomers remain unclear, several biological factors may be implicated. At physiological pH, PFOS exists mostly in its ionic form sulfonate, thus limiting passive diffusion. Active transporters in enterohepatic circulation, such as sodium/taurocholate co-transporting polypeptides (NTCPs) and organic anion transporting polypeptides (OATPs), are shown to mediate the transport of PFOS with species specificity using *in vitro* experiments and computational models.^41–44^ Consequently, The isomer-specific, zonated distribution of PFOS is possibly linked to the zonated distribution and altered expression of relevant hepatic active transporters. Zonation of hepatic OATP transporters has shown a pericentral bias of uptake transporters OATP1A1 and OATP1B2, which is consistent with the higher PFOS levels in the centrilobular region.^45,46^ Interestingly, OAT2, an organic anion transporter with periportal bias, is not believed to play any appreciable role in PFAS transport, further supporting the possible role of OATPs in zonated PFOS distribution.^42,44^ Although evidence so far only showed differential binding affinity of linear vs branched PFOS to serum proteins,^10^ these differences may extend to hepatic transporters and other binding proteins to explain the differential distribution of linear and branched PFOS in hepatic zonation. Importantly, PFOS is reported to alter the expression of its own transporters, as well as many other genes.^35^ FABP (fatty acid-binding protein) has been suggested to be an important binding protein for PFAS *in vitro*,^47^ however *in vivo* studies suggest a minimal role.^48^ FABP is reported to be primarily localized in the periportal region,^49^ consistent with it not significantly sequestering any PFOS isomers. Future work will extend gene expression analysis and explore molecular docking of linear vs branched PFOS with hepatic transporters to further elucidate the mechanism and model the spatially resolved toxicokinetics for linear and branched PFOS.

Our results also for the first time reported PFOS-induced, spatially resolved lipid expression change, characterized by changes in both zonal abundance and zonal distribution. While the no adverse effect level (NOAEL) for hepatic effects is estimated at ∼0.36 mg/kg/day in monkey and rat,^25,50,51^ we see evidence of spatial lipid remodeling even at lower doses, before the emergence of a clear phenotype. We found that lipid zonal distribution, calculated by taking ratios of pericentral and periportal intensities, may serve as a sensitive and unique endpoint to characterize PFOS-induced lipid disruption in addition to abundance analysis. Particularly, calculating zonal distribution within each section can internally normalize the intrinsic variations of MALDI analysis and thus reduce batch effects between different tissues and experimental runs.

Changes in lipid profile demonstrate spatially resolved disruption of lipid homeostasis following PFOS exposure. Lipid changes shared by both zonal abundance and distribution analysis contained what appears to be polyunsaturated fatty acids, such as docosahexaenoic acid (DHA; 22:6), an n-3 polyunsaturated fatty acid that contribute to the regulation of lipid metabolism.^52^ Interestingly, DHA activates the peroxisome proliferator-activated receptor α (PPARα),^53^ a reported target for PFOS exposure and molecular initiating event associated with hepatic steatosis.^54–56^ Furthermore, oxidative stress mediated by hepatic acyl-CoA oxidase 1 (ACOX1), which is regulated by PPARα, was reported as a key event with PFOS-induced alteration in hepatic lipid metabolism, while DHA is a compound that can protect cells from oxidative stress.^56,57^ A decrease of DHA is also common in steatotic and cirrhotic livers.^58^ Arachidonic acid (AA; 20:4)-containing lipid changes suggests a potential increase in the Land’s cycle which has been linked to hepatic steatosis.^59^ AA is also a precursor for eicosanoids implicated in immune responses. Further studies are needed to identify the mechanisms driving lipidomic changes.

### Conclusions and Limitations

MALDI-MS imaging is a powerful imaging modality for the study of endogenous biomolecules and xenobiotics within a spatial context. When coupled with TIMS, the resolution and visualization of isomeric species can provide greater insight into biomolecule profiles and xenobiotic distribution across cell types. Through MALDI-TIMS-TOF MS imaging, we firstly demonstrated that PFOS isomers distribution differently between pericentral and periportal zones, pinpointing the possibility of PFOS isomers’ differential binding capability to hepatic transporters across zones. Spatial lipid perturbations were observed in a zonation-specific manner from chronic exposure at both a high dose (1 mg/kg/day) known to elicit toxicity, and a low dose (0.1 mg/kg/day) treatment comparable to occupationally relevant concentrations in humans.^60^ The multi-modal, spatially resolved analytical workflow for mapping xenobiotic isomer and concomitant disruption in endogenous metabolism has successfully revealed that PFOS isomer structure governs their hepatic zonal distribution and zonation-specific lipid dysregulation, providing key mechanistic insights into PFOS-liver interactions and a powerful toolkit for spatial toxicology research of other xenobiotics.

We acknowledge limitations of this study and suggest future research directions. Our current method parameters did not sufficiently map free fatty acids, preventing deeper investigations into phospholipid biosynthesis and metabolism. Additionally, we focused on analysis of 56 negative-mode lipids that were subjected to iprm-PASEF, which was a subset of all observed signals. Future work will focus on adding more MALDI-MSI modalities with adjacent sections to achieve more comprehensive spatial lipidomics to fully understand PFOS-induced dyslipidemia.

Delocalization was observed in particularly high-dose groups. While the effect is associated with doses, further work should explore optimization of sample preparation for liver samples with higher lipid contents and apply techniques such as sublimation to reduce analyte delocalization.

While revealing on-tissue linear and branched PFOS distribution, this work did not establish absolute quantification of these isomers on tissue. Therefore, it is uncertain whether the level of linear and branched PFOS were present at levels sufficient to contribute to the spatial lipidome changes. Quantification efforts should be implemented for enhanced interpretation of toxicological implications. Future work will include quantitative MSI to quantify region-specific levels of PFOS isomers by performing on-tissue calibration curves or using isotopically labeled standards, and use liquid chromatography (LC)-MS, particularly coupled with ion mobility, to acquire individual isomer concentrations in liver tissues.^61^

## Experimental Methods

### Animal exposure

Male C57BL/6J mice aged postnatal day (PND) 25 were received from the Jackson Laboratory (Bar Harbor, ME) and randomly assigned in groups of 3 to Innocages (Innovive, San Diego, CA) with ALPHA-dri bedding (Shepherd Specialty Papers, Chicago, IL). Mice were housed at 30-40% humidity with a 12-h light/dark cycle. On PND 28 mice were randomly assigned treatment to vehicle control (0.5% Tween-20 water) or PFOS (Accustandard, New Haven, CT; 99% purity; Catalog: PFOS-002N-10MG; lot: 33880) in 0.5% Tween-20 water at concentrations to achieve doses of 0.1 and 1 mg/kg/day. The certificate of analysis notes a 31% representation of branched isomers. PFOS concentrations were recalculated weekly based on body weights and average daily water consumption. Doses reflect the commonly cited no observable adverse effects of 0.1 mg/kg/day,^62,63^ and a higher dose known to elicit hepatic changes. Mice had *ad libitum* access to diet (D09100304; Research Diets Inc., New Brunswick, NJ) and water. Every 4 days, all mice were administered sesame oil by oral gavage as they represent a subset from a larger study. All animal handling activities were performed between Zeitgeber (ZT) times ZT00 and ZT03. On PND 112 (84 days on treatment), between ZT00 and ZT03, mice were euthanized by CO_2_ asphyxiation. Livers were immediately collected, and the left lobe was sectioned for embedding in 5% low-viscosity carboxymethylcellulose (Sigma Aldrich, Cat # C5678). All procedures were approved by the Michigan State University Animal Care and Use Committee (Protocol Number: PROTO202300336) and were guided by the ARRIVE guidelines.^64^

### Materials for MALDI-MSI

LC-MS grade water (Cat # W6212), LC-MS grade acetonitrile (Cat # A955-212), LC-MS grade methanol (Cat # A456-500), Indium-tin oxide (ITO) slides (Cat # NC1365838), and 1,5-diaminonaphthalene (DAN, >98%, Cat # D0101) were purchased from Fisher Scientific (Chicago, IL, U.S.A.). Low viscosity carboxy methylcellulose sodium salt (CMC, Cat # 419273) was purchased from Sigma Aldrich (Milwaukee, WI, U.S.A). Perfluoro-n-octanesulfonate (PFOS, 100 μg/mL, CAT # PFOS-001S) was purchased from AccuStandard (New Haven, CT, U.S.A). Perfluoro-3-methylheptanesulfonate (P3MHpS, 1 μg/mL, CAT # P3MHpS), Perfluoro-4-methylheptanesulfonate (P4MHpS, 1 μg/mL, CAT # P4MHpS), Perfluoro-5-methylheptanesulfonate (P5MHpS, 1 μg/mL, CAT # P5MHpS), Perfluoro-6-methylheptanesulfonate (P6MHpS, 1 μg/mL, CAT # P6MHpS) were purchased from Wellington Laboratories (Overland Park, KS, U.S.A). It should be noted that each standard of P3/P4/P5/P6MHpS also contains perfluoro-3/4/5/6-methylheptanoic acids (P3/P4/P5/P6MHpA) accordingly as part of the purchased standards.

### Sample Preparation for MALDI-MSI

A portion of the left lobe was submerged in 5% w/v CMC and frozen over liquid nitrogen by sitting on a tin-foil boat floating on liquid nitrogen surface. Frozen livers were saved in a -80° C freezer until usage. Frozen embedded tissues were equilibrated at -20° C overnight prior to the day of sectioning. Liver sections were further equilibrated at -10° C for one hour and then sectioned. Livers were mounted on cold chucks using 5% w/v CMC at the base of the frozen tissue mold. 10 µm sections were cut using a cryostat (RWD Minux® FS800) with a chamber temperature of -10° C and chuck temperature of -10° C and were thaw-mounted onto ITO slides. Biological replicates of liver tissues were sectioned and mounted onto ITO slides to generate a layout of randomly paired vehicle control and two treated groups, one section for each, to minimize batch effects among MALDI experimental runs.

Tissues were vacuum sealed and stored at -80° C until ready to image. When ready for analysis, vacuum pouches were taken out of freezer and sat on bench until equilibrated to room temperature, then opened. Slides with tissues are then placed in a vacuum desiccator for 10-20 minutes prior to matrix application and MALDI imaging. Microscopy imaging was performed using a Zeiss Axio Imager Z.2 using brightfield, DAPI, and GFP channels for autofluorescence imaging pre-matrix application and post-MALDI imaging to visualize ablation region on tissue. 1,5-diaminonaphthalene (DAN) was prepared at a concentration of 10 mg/mL in 7:3 acetonitrile/water and applied to tissue sections by an HTX M3+ Sprayer. Pure solvent flow and N_2_ flow were set to 0.100 mL/min and 10 psi, respectively. The nozzle was heated to 55 °C and allowed to pass across the sample 4 times with a track spacing of 2.0 mm. The amount of deposited matrix per unit area was measured as 1.667 μg/mm^2^. Liver samples from 4 animals in control and 5 animals in each treatment were used for MALDI-MSI; one animal from each treated group was later excluded due to their abnormal histopathological features (**Figure S8**), leading to 4 samples per group in final analysis.

### MALDI-TIMS-TOF MS Parameters

MALDI-TIMS-TOF MS imaging was performed by under-sampling 10 μm imaging methods resulting in a 20 μm pixel size subdivided into four 10 μm imaging quadrants. The first quadrant was used to collect quantitative PFOS imaging, the second offset quadrant was used for full positive ion mode lipid imaging, and the third quadrant was used to collect a full negative ion mode lipid profile. We also performed the spotting method to test an aliquot of water with PFOS given to animals to test separation of linear and branched isomers and performed prm-PASEF MS/MS to characterize PFOS isomer structures.

### Negative Ion Mode PFOS Imaging

A timsTOF fleX (Bruker Daltonics, Bremen, Germany) was used to acquire all MALDI imaging data for livers. MALDI parameters in TIMS-mode were optimized to maximize PFOS signal intensity and isomer resolution in the shortest imaging runtime achievable from 100 *m/z* to 1200 *m/z* in negative ionization mode. Mobility and *m/z* calibration were performed through ESI direct infusion using Agilent ESI-L Low Concentration Tune Mix (Cat # AGG1969-85000) purchased from VWR (Radnor, PA, U.S.A). Laser parameters used in data collection include: 10 μm imaging custom laser profile, a laser frequency of 10000 Hz, and 50 laser shots per burst at a laser power of 50%. Ion optics parameters were also tuned, including a Collision Cell Energy of 10.0eV, a Collision RF of 750.0 Vpp, an Ion Transfer Time of 65.0 μs, a Pre Pulse Storage Time of 10 μs, a TIMS Funnel 1 RF of 350.0 Vpp, a TIMS Funnel 2 RF of 250.0 Vpp, a Multipole RF of 250.0 Vpp, a Deflection 1 Delta of −70.0 V, a Δt4 (Accumulation Trap → Funnel 1 In) of −50.0 V, a Δt6 (Ramp Start → Accumulation Exit) of −50.0 V, and a Collision Cell In voltage of −200.0 V. Accumulation time for ions in TIMS funnel was 4.5 ms with a ramp time of 150 ms from 1/K_0_ start value 0.63 V⋅s/cm^2^ to 1/K_0_ end value 1.31 V⋅s/cm^2^.

### Negative Ion Mode Lipid Imaging

Laser parameters used in data collection include: 10 μm imaging laser focusing, a laser frequency of 10000Hz, and 200 laser shots per burst at a laser power of 50%. Ion optics parameters were also tuned, including a Collision Cell Energy of 10.0eV, a Collision RF of 2000.0 Vpp, an Ion Transfer Time of 80.0 μs, a Pre Pulse Storage Time of 10 μs, a TIMS Funnel 1 RF of 450.0 Vpp, a TIMS Funnel 2 RF of 350.0 Vpp, a Multipole RF of 350.0 Vpp, a Deflection 1 Delta of −70.0 V, a Δt4 (Accumulation Trap → Funnel 1 In) of −100.0 V, a Δt6 (Ramp Start → Accumulation Exit) of −50.0 V, and a Collision Cell In voltage of −220.0 V. Accumulation time for ions in TIMS funnel was 20.1 ms with ramp time of 150 ms from 1/K_0_ start value 0.80 V⋅s/cm^2^ to 1/K_0_ end value 1.84 V⋅s/cm^2^. Lipids were collected from a *m/z* window of 150 – 1500 *m/z*. Mobility and *m/z* calibration were performed through ESI direct infusion using Agilent ESI-L Low Concentration Tune Mix.

### Positive Ion Mode Lipid Imaging

Laser parameters used in data collection include: 10 μm imaging laser focusing, a laser frequency of 10000Hz, and 200 laser shots per burst at a laser power of 50%. Ion optics parameters were also tuned, including a Collision Cell Energy of 10.0eV, a Collision RF of 2000.0 Vpp, an Ion Transfer Time of 80.0 μs, a Pre Pulse Storage Time of 10 μs, a TIMS Funnel 1RF of 450.0 Vpp, a TIMS Funnel 2 RF of 350.0 Vpp, a Multipole RF of 350.0 Vpp, a Deflection 1 Delta of 70.0 V, a Δt4 (Accumulation Trap → Funnel 1 In) of 100.0 V, a Δt6 (Ramp Start → Accumulation Exit) of 100.0 V, and a Collision Cell In voltage of 220.0 V. Accumulation time for ions in TIMS funnel was 20.1 ms with ramp time of 150 ms from 1/K_0_ start value 0.80 V⋅s/cm^2^ to 1/K_0_ end value 1.84 V⋅s/cm^2^. Lipids were collected from a *m/z* window of 150 – 1500 *m/z*. Mobility and *m/z* calibration were performed through ESI direct infusion using Agilent ESI-L Low Concentration Tune Mix.

### On-Tissue MS/MS with iprm-PASEF Mode

An ion parallel reaction monitoring-parallel accumulation-serial fragmentation (iprm-PASEF) workflow was applied to annotate lipid species from on-tissue MS/MS using adjacent sections of one replicate of a negative control, and 0.1 mg/kg/day, and 1 mg/kg/day treatment liver. A laser spot size of 20 μm and a raster spacing of 40 μm was used such that up to 4 MS/MS scans per 40 μm pixel could be run per tissue. A precursory MS1 scan was first collected once on a replicate section of a negative control, and 0.1 mg/kg/day, and 1 mg/kg/day treatment liver, followed by 3 MS/MS scans on the same tissue. Adjacent sections of these replicates were immediately subjected to continued MS/MS scans based on initial MS1 feature list. Negative mode iprm-PASEF was performed from a *m/z* window of 150 – 1500 *m/z* and a 1/K_0_ start value 0.85 V⋅s/cm^2^ to 1/K_0_ end value 1.45 V⋅s/cm^2^. Collision cell energies used for fragmentation were defined as 30.00 eV and 50.00 eV for 1/K_0_ 0.80 V⋅s/cm^2^ and 1/K_0_ 1.56 V⋅s/cm^2^, respectively. Collision cell energy was interpolated between these lower and upper limits based on measured 1/K_0_ of analytes. Lipid identification was performed using exact mass matching and MS/MS data against LIPID MAPS database, and using data available in literature.^66^

### PFOS Isomer Characterization by MALDI-TIMS-MS and MS/MS

Mobility/collision cross section (CCS) analysis of PFOS isomers was measured using trapped ion mobility spectrometry (TIMS) at 150 ms and 500 ms ramp times. P3MHpS, P4MHpS, P5MHpS, P6MHpS, n-PFOS standards, and isomeric PFOS content from treatment group water were spotted onto a MALDI target plate and mixed with DAN matrix solution. After crystallization, samples were subject to analysis using a M5 defocused laser profile and the ion optic parameters in the negative ion mode PFOS imaging method. Technical replicates for all spotted samples were collected in n=5. P3MHpS, P4MHpS, P5MHpS, P6MHpS, n-PFOS standards, and isomeric PFOS content from treatment group water and 1 mg/kg/day liver were also measured by MALDI-TIMS-TOF MS/MS via parallel accumulation serial elution fragmentation (PASEF) to characterize isomer structures present in the treatment group water and mouse livers. P3MHpS, P4MHpS, P5MHpS, P6MHpS, n-PFOS standards were spotted onto a MALDI target plate and mixed with DAN matrix solution. After crystallization, samples were subject to analysis using a M5 defocused laser profile and the ion optic parameters in the negative ion mode PFOS imaging method. Technical replicates for iprm-PASEF MS/MS analyzed spotted samples were collected in n=5. Mobility isolation windows for iprm-PASEF MS/MS of standards and treatment water were: n-PFOS (0.810-0.830), P6MHpS (0.804-0.816), P5MHpS (0.795-0.812), P4MHpS (0.793-0.810), P3MHpS (0.792-0.807), treatment water major PFOS isomer (0.810-0.825), and treatment water minor PFOS isomer (0.787-0.803). Fragment ions from technical replicates with a relative abundance of at least 1% to the fragment ion base peak, or S/N > 3.3 were selected for isomer characterization. Fragment ions that were not present in at least 4 of the 5 technical replicates that met the prior intensity criteria were considered not representative of the isomer structure and excluded from averaging. For iprm-PASEF imaging of 1 mg/kg/day PFOS treatment liver, the sample was prepared as described the “Sample Preparation for MSI” section. Ramp times of 0.810-0.824 and 0.786-0.800 were used for the major and minor PFOS isomers, respectively. Fragment ions with a relative abundance of at least 1% to the fragment ion base peak, or S/N > 3.3 were selected for isomer characterization on tissue. The iprm-PASEF MS/MS was performed using a 40 eV collision energy using a ramp time of 500 ms. Other instrument parameters can be found from our previous work.^30^

### Assessing Detection Limits of PFOS in 0.1 mg/kg/day Treatment Livers

MALDI-TIMS-TOF imaging .d files were imported into Compass DataAnalysis (Version 6.1) for detection limit calculations. Three regions of low PFOS intensity were selected randomly for S/N ratio calculations across each imaged 0.1 mg/kg/day livers. Each region was user-defined in DataAnalysis to generate a mobilogram based on isolated pixels using the SCiLS rendered ion image as reference. The major and minor PFOS isomer mobilogram peaks were visualized by applying a global extracted ion mobility (EIM) filtering using the observed PFOS [M-H]^-^ion ± 10 ppm. Mobilogram peak areas defined from sub-regions which contained both major and minor PFOS isomers were integrated for isomer-specific S/N calculations. S/N calculations were performed as described from our previous work. ^30^

### Materials for Immunofluorescence

1 X PBS (Gibco; Catalog # 10010-023), CytoFix/Cytoperm (BD Biosciences; Catalog: 554722), Bovine Serum Album (Sigma; Catalog: A1933-5G), Triton X-100 solution (Sigma; Catalog: 93443-100mL), PAP pen (Sigma; Catalog: Z377821), Hoechst 33342 (Biotium; Catalog: 40046), Fc mouse block CD16/CD32 (BD Biosciences; Catolog: 553141), Alexa Fluor Plus 647 secondary antibody (Invitrogen; Catalog: A32795TR), Primary ASS1 antibody (Abcam; Catalog: AB170952), Glutamine Synthetase Coralite ®PLUS 488-conjugated (Proteintech; Catalog: CL488-66323-2), Fluoromount-G (SouthernBioTech;Catalog: 0100-01).

### Immunofluorescence imaging of zonation markers

Following MALDI imaging, the DAN matrix was washed off by sequentially submerging the slides in 95% ethanol, 70% ethanol, and three washes with deionized, nuclease-free molecular biology grade water. Tissue sections were air dried, and a PAP pen was used to create a hydrophobic barrier around the tissue section. After the barrier dried, the tissue was rehydrated in 1 X PBS for 40 minutes at room temperature then fixed with ice cold CytoFix/Cytoperm solution (1:4) for 20 minutes. Sections were incubated at 37°C for 60 mins, protected from light, in staining buffer (1% BSA, 0.1% Triton X-100, 1X PBS) with Mouse BD Fc Block CD16/CD32 (1:100) and Hoechst 33342 (nuclei stain; 1:1000). The solution was aspirated, then the slides incubated with anti-ASS1 antibody (portal hepatocyte marker; 1:100) and Coralite® PLUS 488 conjugated anti-GLUL (central hepatocyte marker; 1:100) in staining buffer at 37℃ for 60 minutes, protected from light. Slides were rinsed three times with 1 X PBS then incubated with Alexa Fluor Plus 647 secondary antibody (1:100) in staining buffer at 37℃ for 60 minutes, protected from light, followed by another three rinses of 1 X PBS. Coverslips were mounted using Fluoromount G and imaged using a Leica THUNDER microscope at the MSU IQ Microscopy Core Facility (<u>microscopy.iq.msu.edu</u>). QuPath was used for analysis and preparation of representative photomicrographs.^66^

### Data Analysis

All MALDI-TIMS-TOF MS imaging data was visualized and analyzed using SCiLS Lab Version 2025b (Bruker Daltonics, Bremen, Germany). Data was total ion count (TIC) normalized. Segmentation of imaged tissues was performed using the bisecting k-means method and correlation distance metric to unbiasedly outline the tissue region and define zones. Strong denoising was applied to segmentation of imaged livers to definition clarity of zonation features. Pericentral and periportal zonal regions were validated against same tissue immunofluorescence staining of established markers to confirm accuracy of segmentation. Pericentral and periportal regions created from segmentation were used for lipid profile changes and PFOS zonal distribution analysis. An initial feature list from imaging data using the T-ReX^3^ (TIMS) algorithm with a spatial averaging of 2x2, 20% coverage and 0.3% relative intensity threshold was used to generate a comprehensive lipidomic feature table for tentative lipid identification. A second feature list was generated from imaging data using the T-ReX^3^ (TIMS) algorithm with a spatial averaging of 2x2, 10% coverage and 1% relative intensity threshold was created to identify lipids of sufficient intensity for on-tissue iprm-PASEF annotation. A *m/z* interval window of 15 ppm window was used to ensure that peak areas were adequately covered for quantitative purposes. Lipids from segmented pericentral and periportal zones were identified based on the following criteria: the ion has a probable assignment based on MS1 accurate mass matching to LIPID MAPS databases (https://www.lipidmaps.org); and the ion image is clear in quality across replicates. For iprm-PASEF imaging of lipids, an additional criterion was implemented where lipids must have an average TIC-normalized intensity of 10,000 or greater across the tissue area were selected for on-tissue MS/MS annotation to ensure adequate fragmentation signal for reliable annotation. For all lipids, ccs matching was evaluated against existing literature from drift tube ion mobility spectrometry (DTIMS) measurements.^67-69^ Average TIC-normalized intensities of lipids, n-PFOS and br-PFOS from segmented pericentral and periportal zones were exported to a CSV file for data analysis. Segmented pericentral:periportal ratios for lipids, n- and br-PFOS were calculated to represent the pericentral:periportal zonal distribution of lipids and zonal distribution of n- and br-PFOS from the following two equations:

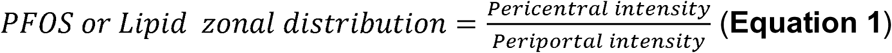

A log_2_ transformation was applied to these ratios such that zero represents equal zonal distribution. A paired student’s t-test was performed on the log_2_-transformed data of PFOS isomer intensities using GraphPad Prism (V. 11.0.0) to determine if n-PFOS and br-PFOS distribute differently between zones in each treatment group. For lipid zonal distribution analysis, Dunnett’s multiple comparisons were used to compare the low-and high-dose groups separately against the control group, and the resulting Dunnett-adjusted p-values were further corrected using the Benjamini-Hochberg false discovery rate (FDR) procedure to correct multiple comparisons across lipid species. Statistical significance was defined as FDR-adjusted q < 0.05.

To determine zone-specific changes in lipid profiles, fold change (FC) between treated and control group were calculated in pericentral and periportal zones separately:

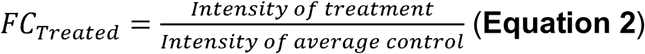

A log_2_ transformation was applied to FC of lipids prior to statistical analysis such that zero represents no change between treated and control. Statistical comparisons were performed on *log*_₂_*FC* values using one-way ANOVA followed by a Dunnett multiple comparison test to compare treatment values to the control values. Resulting differences were interpreted and presented as log_2_(FC) for interpretability. Statistical significance for all analyses was defined as p < 0.05.

For hierarchical clustering of zonal distribution, lipid features were standardized across samples by row-wise Z-scoring. Sample clustering was performed using Euclidean distance with Ward.D2 linkage, while lipid clustering was performed using Pearson correlation distance (1 - r) with average linkage. Clustering was first conducted in all treatment groups, and a secondary analysis for only control and low-dose groups was also performed using the same settings.

To evaluate within-sample zonal differences while accounting for spatial autocorrelation among neighboring pixels, conditional autoregressive (CAR) model was fitted following the framework provided by Cassess et al. For each liver section, the response was the pixel-wise log2-transformed linear:branched PFOS intensity ratio, with hepatic zonation included as a fixed effect. Spatial neighborhoods were defined using Euclidean distances of 1 to 4 pixels. The numerically stable model with the lowest AIC was retained. Zona effect estimates, fold changes, and p-values were reported. Residue Moran’s I was used to assess remaining spatial autocorrelation. Hierarchical clustering and CAR were both performed and results plotted using R. ChatGPT-5.6 Sol (OpenAI) was used to assist with generation of R code for hierarchical clustering and CAR modeling. All generated codes were reviewed, modified as needed, and executed by the authors.

## Supporting Information

Analysis of isomeric PFOS using MALDI-TIMS-MS/MS, vehicle control ion images, immunofluorescence example, multi-modal results on 1 mg/kg/day PFOS-exposed liver, example segmentation and overlaying with immunofluorescence, representative photomicrographs of livers with abnormal pathology, signal-to-noise ratio of PFOS isomers on tissue, complete heatmaps of lipid zonal abundance and distribution.

## Author Contributions

A.J.R.: conceptualization, methodology, validation, formal analysis, investigation, data curation, visualization, writing – original draft, writing – review & editing; J.A.M.: methodology, formal analysis, investigation, data curation, visualization, writing – review & editing; S.S.: methodology, writing – review & editing; R.N.: conceptualization, methodology, resources, data curation, writing – review & editing, supervision, project administration, funding acquisition; T.A.Q.: conceptualization, methodology, resources, data curation, writing – original draft, writing – review & editing, supervision, project administration, funding acquisition.

## Conflict of Interest

The authors declare that there are no conflicts of interest.

## Data Availability Statement

The data that support the findings of this study are available in the Methods, Results, and Supplemental Information of this article. Raw MSI and post-MALDI data are deposited in the MassIVE (MSV000102850) and BioImage Archive (S-BIAD3956) repositories.

## Acknowledgements

This work was supported by startup funding from Michigan State University to T.A.Q. and R.N., and a Starter Grant from the Society for Analytical Chemists of Pittsburgh and a Research Award from the American Society of Mass Spectrometry to T.A.Q. This work was supported in part by USEPA STAR Program (No. 84095601). This document has not been formally reviewed by the funding agency. The views expressed in this document are solely from the authors and do not necessarily reflect those from the funding agency. The agency does not endorse any products or commercial services mentioned in this paper.

## Notes

### Competing Interest Statement

The authors have declared no competing interest.

### Summary of Updates

This manuscript has been significantly revised to add: - Increased biological replicates - Quantitative analysis of PFOS zonal ratios between pericentral and periportal for both linear and branched PFOS - Characterization of branched PFOS using authentic standards with TIMS and iprm-PASEF - In-depth characterization of spatial lipidome in control and exposed samples with on-tissue MS/MS - Quantitative analysis of lipid zonal abundance and zonal ratios between PC/PP regions.

