## Supplemental Information for "Multi-Modal Mass Spectrometry Imaging Reveals Isomer-Specific PFOS Distribution and Spatial Lipid Disruption in Mouse Liver"

**Table of Contents**

**Supplemental Texts**. Analysis of isomeric PFOS using MALDI-TIMS-MS/MS.

**Table S1.** Observed *m/z* and collision cross sections (CCS) values for PFOS isomers.

**Table S2.** Average relative abundance of PFOS isomer product ions from iprm-PASEF.

**Figure S1.** Analysis of PFOS isomers by trapped ion mobility spectrometry (TIMS).

**Figure S2. i**prm-PASEF MS/MS spectra of PFOS isomers.

**Figure S3.** Detection and spatial distribution of PFOS on exposed liver tissue sections

**Figure S4.** Vehicle control ion images show no PFOS present in livers.

**Figure S5:** Comprehensive PFOS ion images from all imaged livers in low dose (0.1 mg/kg/day) and high dose (1 mg/kg/day) treatment groups.

**Figure S6**. Post-MALDI immunofluorescence imaging of zonal hepatocyte markers.

**Figure S7:** Multi-modal imaging results on the same 1 mg/kg/day PFOS-exposed liver.

**Figure S8.** Example segmentation analysis of a vehicle tissue towards accurate pericentral and periportal zonation.

**Figure S9**. Example of validating segmented regions against immunofluorescence

**Figure S10**. Representative photomicrographs of livers of PFOS treated mice

**Figure S11.** Signal-to-noise ratio of PFOS isomers across low intensity liver regions.

**Figure S12.** On-tissue ionization efficiency of PFOS isomers by MALDI-TIMS-MSI.

**Figure S13**. Spatial analysis of PFOS zonal distributions for low-dose groups (0.1 mg/kg/day) using conditional autoregressive (CAR) modeling

**Figure S14**. Spatial analysis of PFOS zonal distributions for high-dose groups (1 mg/kg/day) using conditional autoregressive (CAR) modeling.

**Figure S15.** Average spectra of vehicle control tissues from negative mode and positive mode imaging.

**Figure S16.** Positive mode lipid imaging visualizing a biomarkers of periportal and pericentral zones in vehicle control livers.

**Table S3**. MS/MS annotations from iprm-PASEF analysis.

**Figure S17:** Heatmaps illustrating individual replicate response and lipid abundance fold change to PFOS treatment.

**Figure S18**. Heatmap of zonal ratios for lipids and treatment groups.

**Figure S19**. Hierarchical clustering with control and low-dose group only.

**Figure S20:** Comprehensive lipid ion images showing lipids which undergo statistically significant zonal localization indicated by Figure 5C.

**Analysis of isomeric PFOS using MALDI-TIMS-MS and MS/MS**

Isomeric PFOS content from treatment group water was directly measured by MALDI-TIMS-TOF MS/MS to identify potential isomers present in the purchased standards used in treatment studies. Trapped ion mobility spectrometry (TIMS) ramp times ranging from 73 ms to 800 ms over a mobility window of 0.63-1.31 1/K_0_ were investigated to optimize PFOS isomer separation in treatment water samples. From this ramp time optimization, it was found that a minimum ramp time of 150 ms was required to resolve the two isomer peaks enough such that software could identify and annotate both PFOS isomers present in treatment studies. Baseline resolution was not achieved by using this 150 ms ramp time, or by extending ramp times used for isomer separation in water samples. Analysis of several PFOS isomers including linear PFOS (n-PFOS), perfluoro-6-methylheptanesulfonate (P6MHpS), perfluoro-5-methylheptanesulfonate (P5MHpS), perfluoro-4-methylheptanesulfonate (P4MHpS), and perfluoro-3-methylheptanesulfonate (P3MHpS) confirmed that at moderate ramp times (150 ms) and extended ramp times (500 ms) baseline resolution could not be achieved between standards of equivalent concentrations **(Figure S1A-B)**. Notably, TIMS could not resolve P3MHpS, P4MHpS, and P5MHpS isomers from one another. Measured collision cross section (CCS) values from TIMS confirmed that at both 150 ms and 500 ms ramp times, P3MHpS, P4MHpS, and P5MHpS were too structurally similar to differentiate with ion mobility alone **(Table S1)**. This limitation has been observed in other ion mobility modalities, complicating isomer identification in complex mixtures.^1^ Analysis of treatment water and treatment liver mobilograms against analytical standards strongly suggested that the major PFOS isomer (Ma-PFOS) corresponded to n-PFOS whereas the minor PFOS isomer (Mi-PFOS) corresponded to a potential branched mixture of P3MHpS, P4MHpS, P5MHpS, and potentially other branched isomers **(Figure S1C-F)**. This led to the usage of a 150 ms ramp time as the optimal setting to balance isomeric separations and imaging run speeds.

To better characterize the branched isomer mixture present in treatment water and liver samples, ion parallel reaction monitoring – parallel accumulation serial fragmentation (iprm-PASEF) was utilized for isomer-resolved MS/MS data. MS/MS spectra of isomer standards showed differences in fragment ion intensities **(Figure S2)**, however, these intensity differences could not solely be used for compound identification in complex mixtures. Further examination of MS/MS spectra for isomer-specific diagnostic fragments found few diagnostic fragments that could be used for isomer identification in samples. Among the tested PFOS isomers, a potential diagnostic fragment of *m/z* 379.9376 was discovered, which corresponds to the molecular formula of [C_6_F_12_SO_3_]^-^ **(Table S2).** While this fragment was also identified in n-PFOS, our n-PFOS analytical standard displayed branched isomer impurities from which this fragment ion could originate from. To the best of our knowledge, this fragment ion has not been reported in n-PFOS,^2^ and for it to see use as a diagnostic fragment for P5MHpS, baseline resolution between isomers should be achieved to clearly attribute fragment ion origin to one isomer. P4MHpS yielded the diagnostic fragment *m/z* 310.9429 which corresponds to the molecular formula of [C_5_F_9_SO_3_]^-^ **(Table S2)**. This fragment ion was present at low relative abundances suggesting it was difficult to form or an unstable fragment ion. P3MHpS yielded two different diagnostic fragment ions in *m/z* 318.9376 and *m/z* 260.9456 which correspond to the molecular formulas [C_5_S_10_SO_3_]^-^ and [C_4_F_7_SO_3_]^-^, respectively **(Table S2)**. Interestingly, while P3MHpS, P4MHpS, and potentially P5MHpS/n-PFOS possessed diagnostic fragment ions, P6MHpS did not yield its own diagnostic fragment ion. This prevented confirming its presence in the treatment water and liver samples.

Comparison of Ma-PFOS and Mi-PFOS spectra from treatment water showed some discrepancies between individual isomer standards. Relative intensities differed in both the Ma-PFOS and Mi-PFOS MS/MS spectra likely due to intensity convolution from a mixture of isomers **(Figure S2)**. Diagnostic fragments for both P3MHpS and P4MHpS were found to be missing from MS/MS data **(Table S2)**. When preparing samples for mobility and MS/MS analysis, standards were prepared at a concentration of 500 ng/mL whereas treatment water samples were prepared at a concentration of 2 μg/mL. Despite the treatment water having a higher concentration, intensities were found to be lower than those of the analytical standards. We suspect the presence of a nonionic surfactant in tween-20 at a concentration of 0.5% is introducing ion suppression in the water samples, which may compromise detection of low-abundance fragment ions. When looking at the MS/MS spectra of 1 mg/kg/day livers, we observed significantly clearer MS/MS data allowing for more accurate comparisons against standards. Ma-PFOS present in livers showed similar MS/MS fragment ions and relative intensities to n-PFOS **(Table S2, Figure S2)**, and near identical CCS values **(Table S1),** confirming that the major isomer present in treatment is n-PFOS. Contrary to the Ma-PFOS MS/MS spectra, the Mi-PFOS spectra did not match any single PFOS isomer standard, confirming that the Mi-PFOS peak is a complex mixture of unknown isomers. In agreement with CCS values indicating that the Mi-PFOS consisted of P3MHpS and/or P4MHpS **(Table S1)**, diagnostic fragments for both P3MHpS (*m/z* 318.9376 and *m/z* 260.9456) and P4MHpS (*m/z* 310.9429) were identified in the Mi-PFOS mixture **(Table S2)**. However, without fully establishing *m/z* 379.9376 as a unique diagnostic fragment for P5MHpS, we cannot rule out the potential presence of P5MHpS present in the Mi-PFOS isomer mixture.

Beyond known fragment ions from tested isomer standards, a new fragment ion not present in tested isomers appeared in the treatment water, and was fully resolved to the Mi-PFOS peak in liver samples. A major diagnostic peak was identified as 418.9728 *m/z* which corresponds to the neutral loss of the sulfonate head group [M-SO_3_H]^-^ **(Table S2, Figure S2)**. MS/MS data available for isomeric perfluorosulfonic acids (PFSAs) and their derivatives show that the production of the [M-SO_3_H]^-^ ion or [M-head group]^-^ is a distinct diagnostic fragment for 1-methyl substituted branched isomers, ^3–5^ suggesting that a perfluoro-1-methylheptanesulfonate (P1MHpS) or another unresolved PFOS isomer may be present in the branched mixture which generate [M-SO_3_H]^-^. However, with the absence of direct confirmation with authentic standards, retention time from liquid chromatography, or further resolved ion mobility separation, we cannot confirm its presence. Future work should seek LC-MS coupled with ion mobility for its detection in liver samples.

**Table S1.** Observed *m/z* and collision cross sections (CCS) values for PFOS isomers

|  | 150 ms Ramp Time | | | | | 500 ms Ramp Time | | | | |
| --- | --- | --- | --- | --- | --- | --- | --- | --- | --- | --- |
|  | **n-PFOS** | | | | | **n-PFOS** | | | | |
|  | Replicate 1 | Replicate 2 | Replicate 3 | Replicate 4 | Replicate 5 | Replicate 1 | Replicate 2 | Replicate 3 | Replicate 4 | Replicate 5 |
| *m/z* | 498.9286 | 498.929 | 498.9293 | 498.9294 | 498.9286 | 498.9298 | 498.9292 | 498.9298 | 498.9296 | 498.9293 |
| CCS (Å^2^) | **168.3** | **168.6** | **168.6** | **168.4** | **168.5** | **167.9** | **168.0** | **168.0** | **168.0** | **168.0** |
|  | **P6MHpS** | | | | | **P6MHpS** | | | | |
|  | Replicate 1 | Replicate 2 | Replicate 3 | Replicate 4 | Replicate 5 | Replicate 1 | Replicate 2 | Replicate 3 | Replicate 4 | Replicate 5 |
| *m/z* | 498.9303 | 498.9302 | 498.9288 | 498.9296 | 498.9284 | 498.9287 | 498.9292 | 498.9295 | 498.9296 | 498.9294 |
| CCS (Å^2^) | **166.4** | **166.4** | **166.5** | **166.5** | **166.6** | **166.0** | **166.1** | **166.1** | **166.1** | **166.4** |
|  | **P5MHpS** | | | | | **P5MHpS** | | | | |
|  | Replicate 1 | Replicate 2 | Replicate 3 | Replicate 4 | Replicate 5 | Replicate 1 | Replicate 2 | Replicate 3 | Replicate 4 | Replicate 5 |
| *m/z* | 498.9285 | 498.9292 | 498.9299 | 498.9289 | 498.9294 | 498.929 | 498.9298 | 498.9294 | 498.9297 | 498.9298 |
| CCS (Å^2^) | **165.2** | **165.1** | **165.0** | **165.2** | **165.1** | **164.6** | **164.6** | **164.5** | **164.5** | **164.7** |
|  | **P4MHpS** | | | | | **P4MHpS** | | | | |
|  | Replicate 1 | Replicate 2 | Replicate 3 | Replicate 4 | Replicate 5 | Replicate 1 | Replicate 2 | Replicate 3 | Replicate 4 | Replicate 5 |
| *m/z* | 498.9297 | 498.9301 | 498.9301 | 498.9292 | 498.9298 | 498.9297 | 498.9297 | 498.9301 | 498.9301 | 498.93 |
| CCS (Å^2^) | **164.5** | **164.5** | **164.7** | **164.6** | **164.6** | **164.0** | **164.2** | **164.1** | **164.1** | **164.3** |
|  | **P3MHpS** | | | | | **P3MHpS** | | | | |
|  | Replicate 1 | Replicate 2 | Replicate 3 | Replicate 4 | Replicate 5 | Replicate 1 | Replicate 2 | Replicate 3 | Replicate 4 | Replicate 5 |
| *m/z* | 498.9297 | 498.9291 | 498.9287 | 498.9293 | 498.9288 | 498.9298 | 498.93 | 498.9297 | 498.9295 | 498.93 |
| CCS (Å^2^) | **164.3** | **164.5** | **164.4** | **164.4** | **164.3** | **164.0** | **164.1** | **163.9** | **164.1** | **164.0** |
|  | **Observed Major PFOS Isomer in Treatment Water** | | | | | **Observed Major PFOS Isomer in Treatment Water** | | | | |
|  | Replicate 1 | Replicate 2 | Replicate 3 | Replicate 4 | Replicate 5 | Replicate 1 | Replicate 2 | Replicate 3 | Replicate 4 | Replicate 5 |
| *m/z* | 498.9301 | 498.9296 | 498.9297 | 498.9293 | 498.9293 | 498.9302 | 498.929 | 498.93 | 498.9293 | 498.9295 |
| CCS (Å^2^) | **168.1** | **168.0** | **168.3** | **168.1** | **168.2** | **167.7** | **167.7** | **167.4** | **167.7** | **167.6** |
|  | **Observed Minor PFOS Isomer in Treatment Water** | | | | | **Observed Minor PFOS Isomer in Treatment Water** | | | | |
|  | Replicate 1 | Replicate 2 | Replicate 3 | Replicate 4 | Replicate 5 | Replicate 1 | Replicate 2 | Replicate 3 | Replicate 4 | Replicate 5 |
| *m/z* | 498.9297 | 498.9292 | 498.9302 | 498.9299 | 498.9296 | 498.9295 | 498.301 | 498.9284 | 498.9304 | 498.9296 |
| CCS (Å^2^) | **164.1** | **163.5** | **165.1** | **164.2** | **165.3** | **163.9** | **164** | **163.7** | **163.6** | **164** |

**Table S2. Average relative abundance (≥1%) of PFOS isomer product ions from iprm-PASEF MS/MS (40 eV CE, 500ms ramp time).** Most abundant fragment ion was established as base peak (100%).

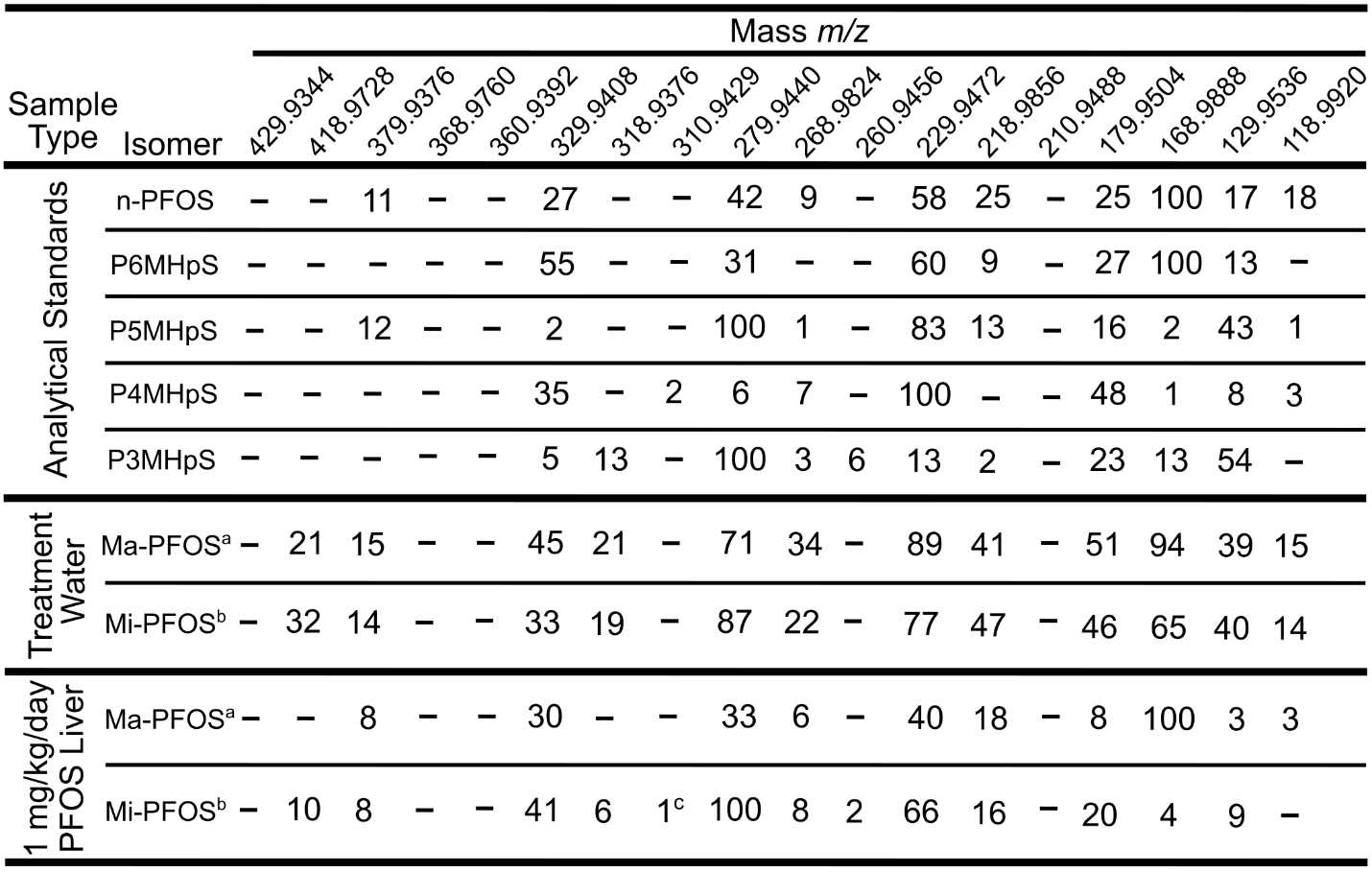

^a^Major unknown PFOS isomer peak abbreviated as Ma-PFOS.

^b^Minor unknown PFOS isomer peak abbreviated as Mi-PFOS.

^c^Relative abundance measured as 0.85% with S/N>3.3 was rounded to 1% and included for isomer identification.

—: not detected in at least 4 of 5 technical replicates.

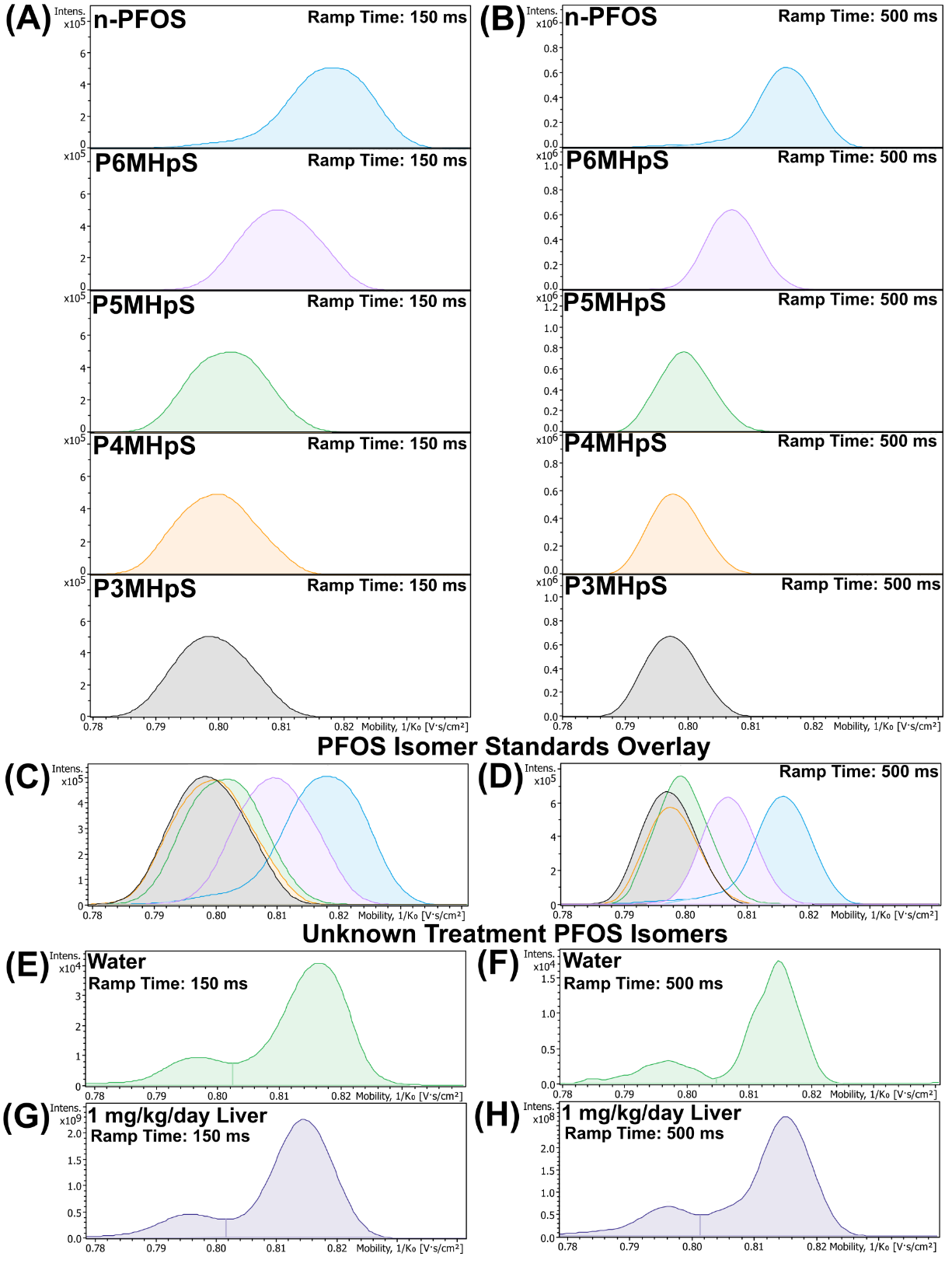

**Figure S1.** **Analysis of PFOS isomers by trapped ion mobility spectrometry (TIMS).** (A-B) Mobilograms of n-PFOS, P6MHpS, P5MHpS, P4MHpS, and P3MHpS using a 150 ms and 500 ms ramp time. (C-D) Overlays of individually measured PFOS isomer mobilograms at 150 ms and 500 ms ramp times. (E-F) Mobilogram of a water aliquot from 1 mg/kg/day treatment group showing isomers present. (G-H) Mobilogram of PFOS extracted from MALDI-TIMS-TOF mass spectrometry imaging of a 1 mg/kg/day liver sample at 150 ms and 500 ms ramp times.

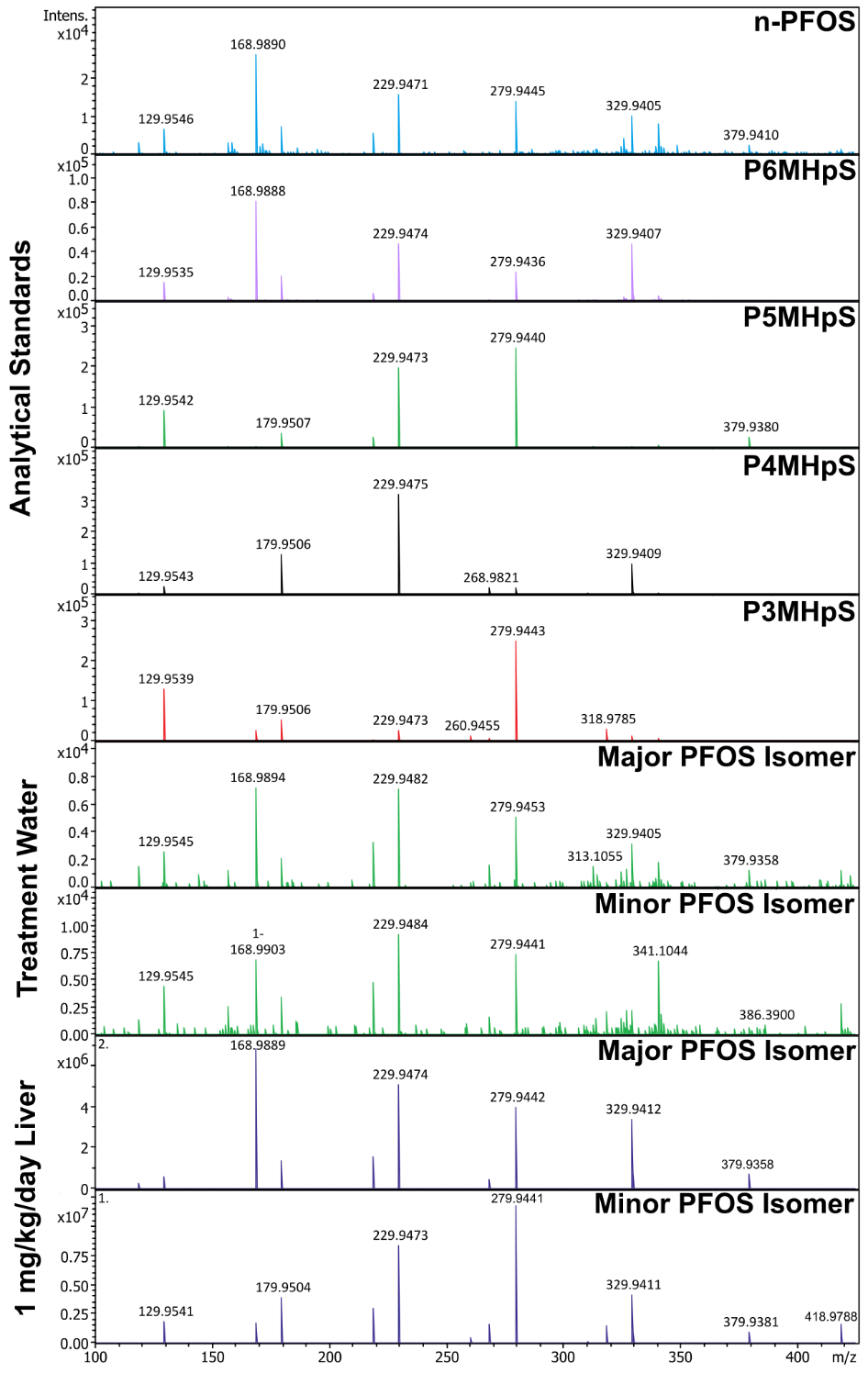

**Figure S2. iprm-PASEF MS/MS spectra of PFOS isomers (40 eV CE, 500ms ramp time).** MS/MS spectra of PFOS isomers present in analytical standards, treatment water, and 1 mg/kg/day liver reveal differences in product ion and ratios.

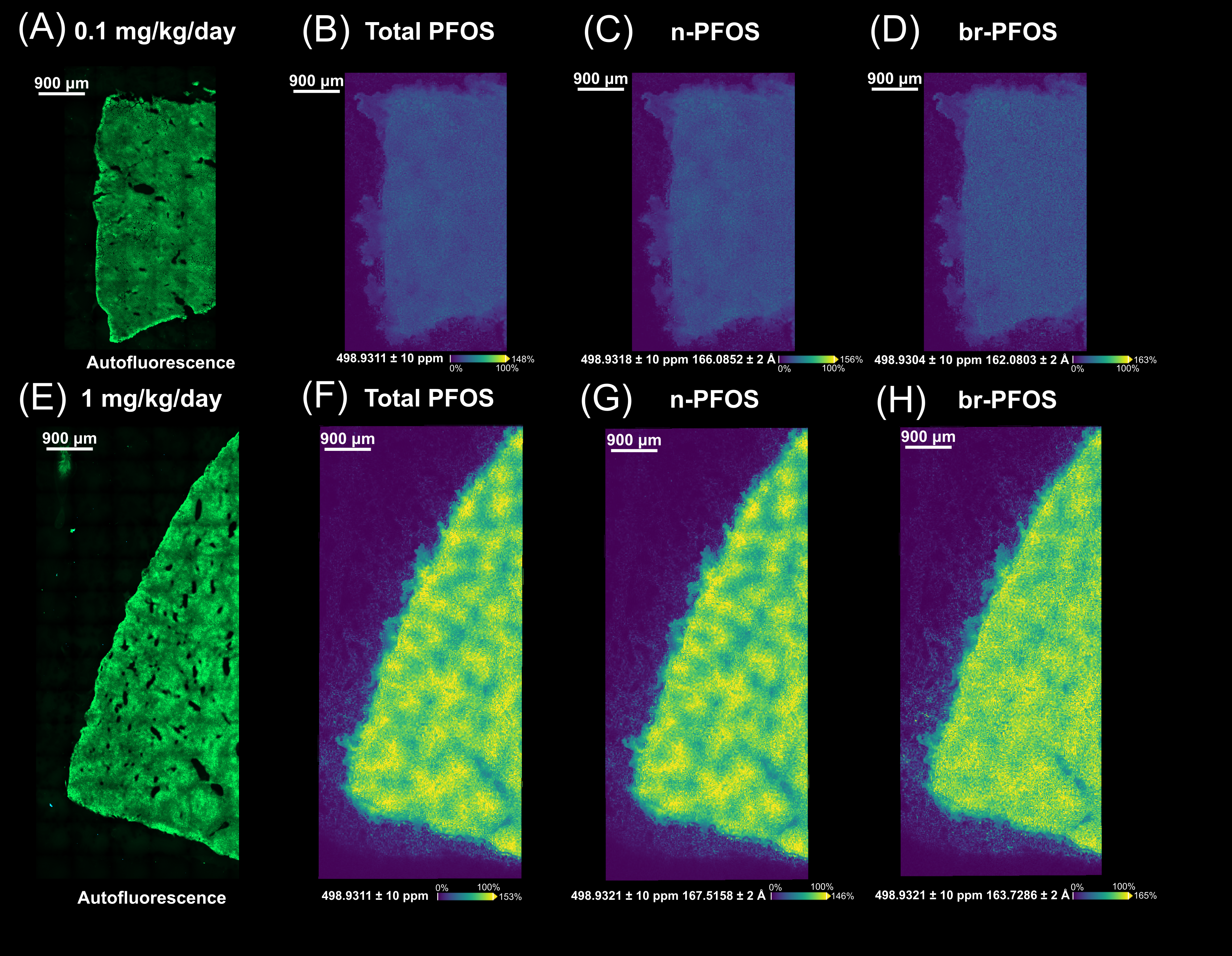

**Figure S3.** Detection and spatial distribution of PFOS on exposed liver tissue sections (A-D) Pre-MALDI brightfield and green autofluorescence providing tissue area overview and MALDI-MSI ion images for total, linear, and branched PFOS from 0.1 mg/kg/day PFOS-exposed liver samples. (E-H) Images from 1 mg/kg/day PFOS-exposed liver samples. Intensity scaling is not adjusted to visualize contrast of PFOS accumulation differences between treatment groups. MALDI-MSI parameters: matrix-DAN; polarity-negative; lateral spatial resolution-20 μm; other parameters-see Experimental methods.

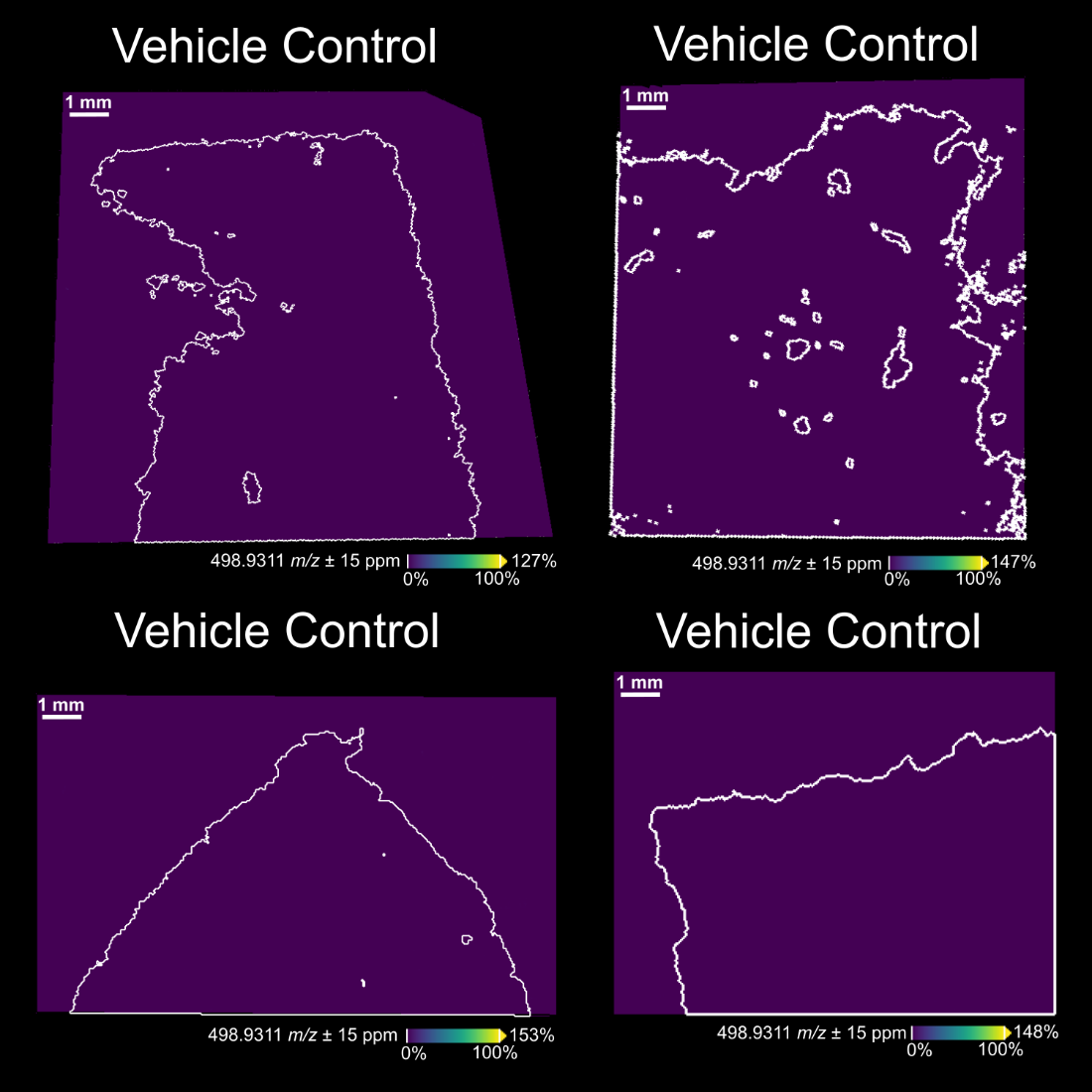

**Figure S4. Vehicle control ion images show no PFOS present in livers.** PFOS was not detected by MALDI-TIMS-TOF-MSI from vehicle controls indicating that food was not a major source of incidental PFOS exposure, and that sample preparation did not introduce cross-contamination.

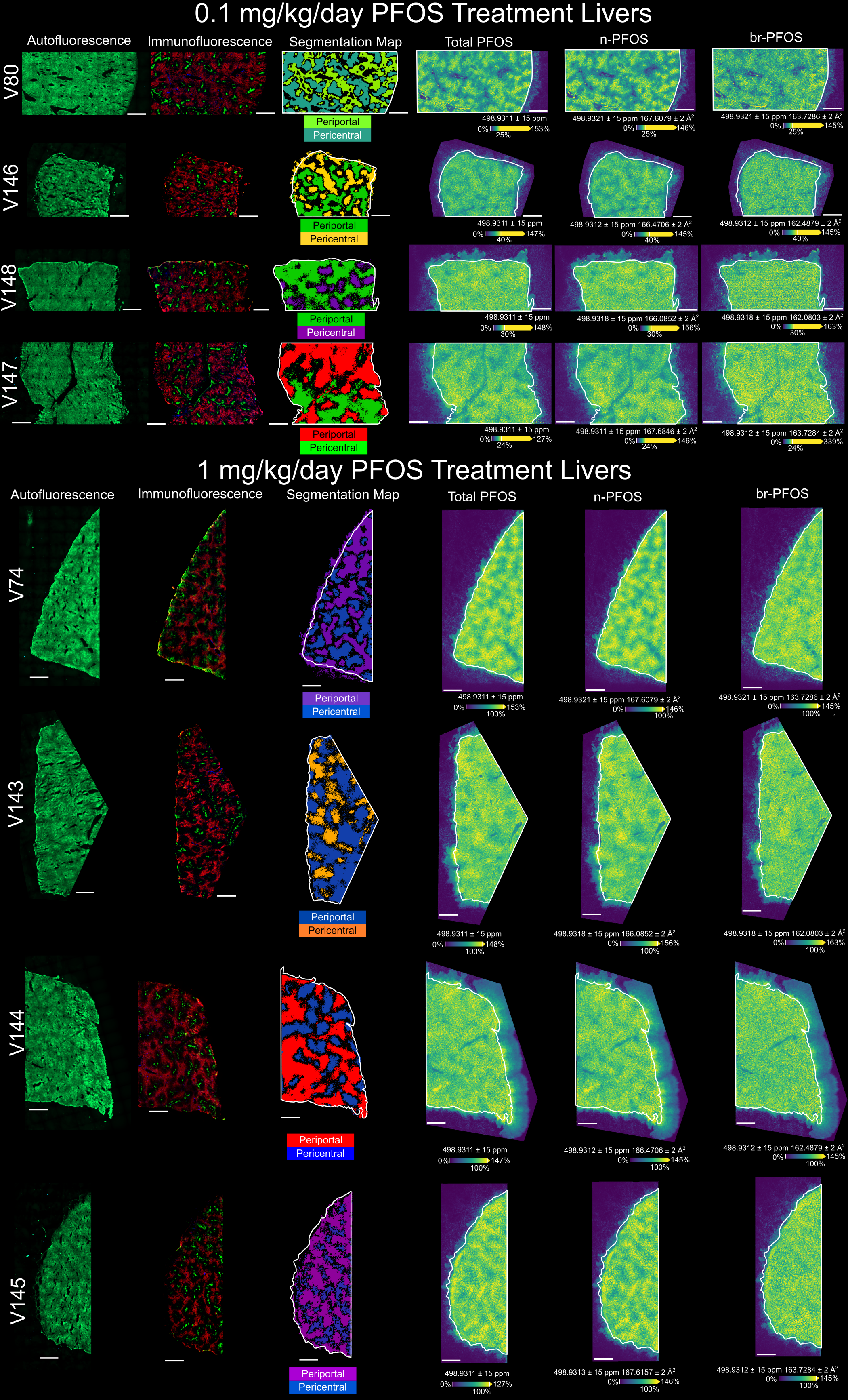

**Figure S5:** Comprehensive PFOS ion images from all imaged livers in low dose (0.1 mg/kg/day) and high dose (1 mg/kg/day) treatment groups. Tissue outlines are indicated by the white bordering around ion images. Intensity thresholds are scaled in 0.1 mg/kg/day samples to provide similar contrast for localization comparisons against 1 mg/kg/day livers. Scale bars represent 900 μm.

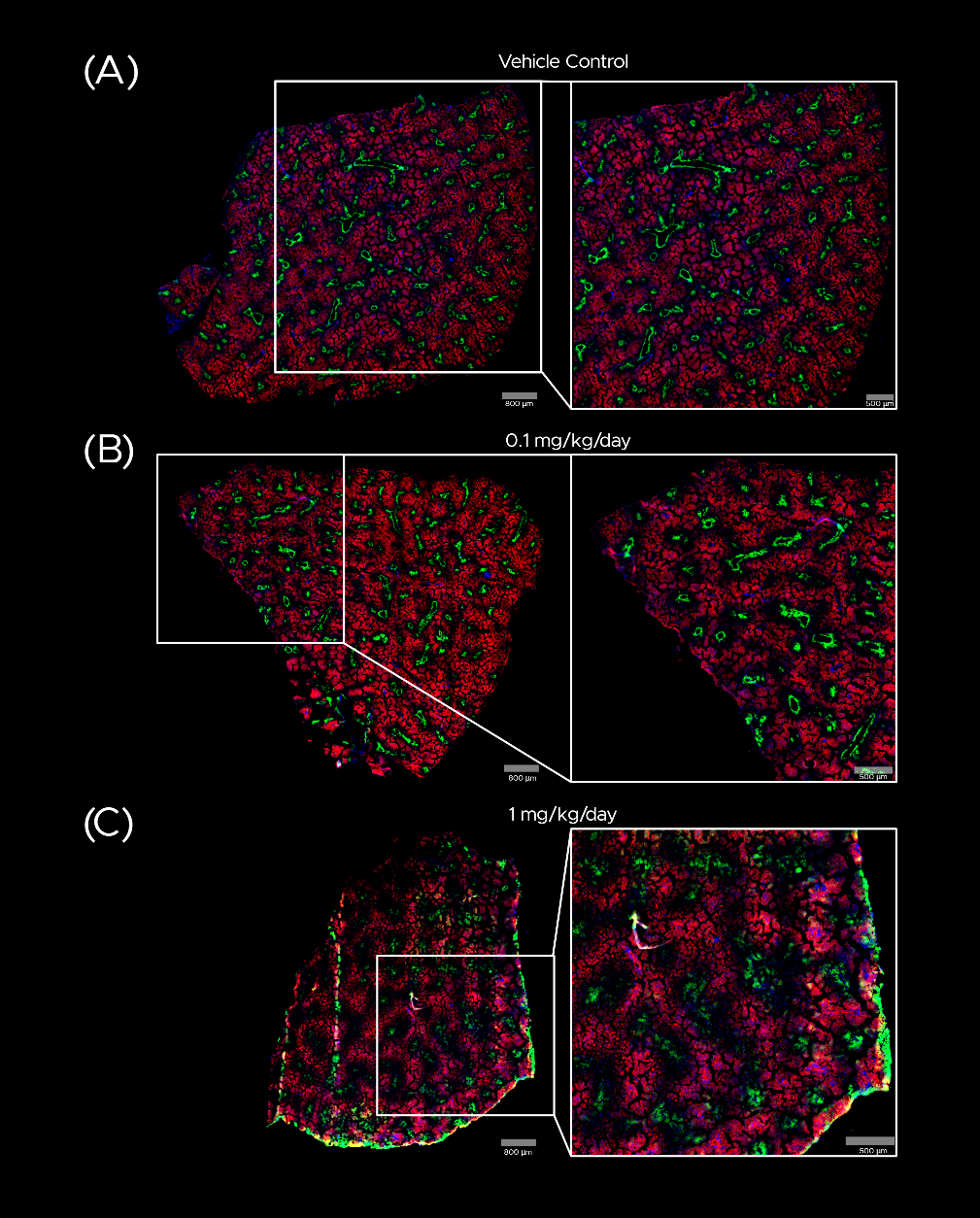

**Figure S6. Immunofluorescence imaging of zonal hepatocyte markers of MALDI imaging liver sections.** Representative photomicrographs are shown for livers from (A) vehicle control, (B) 0.1 mg/kg/day PFOS, and (C) 1 mg/kg/day PFOS treated mice. Full tissue image shown on the left and a magnified portion representing the MALDI imaged region is shown on the right. Livers were stained for Hoechst 33342 (Nuclei; blue), ASS1 (portal hepatocytes; red), and GLUL (central hepatocytes; green). Scale bar represents 800 µm and 500 µm for full size and magnified sections, respectively.

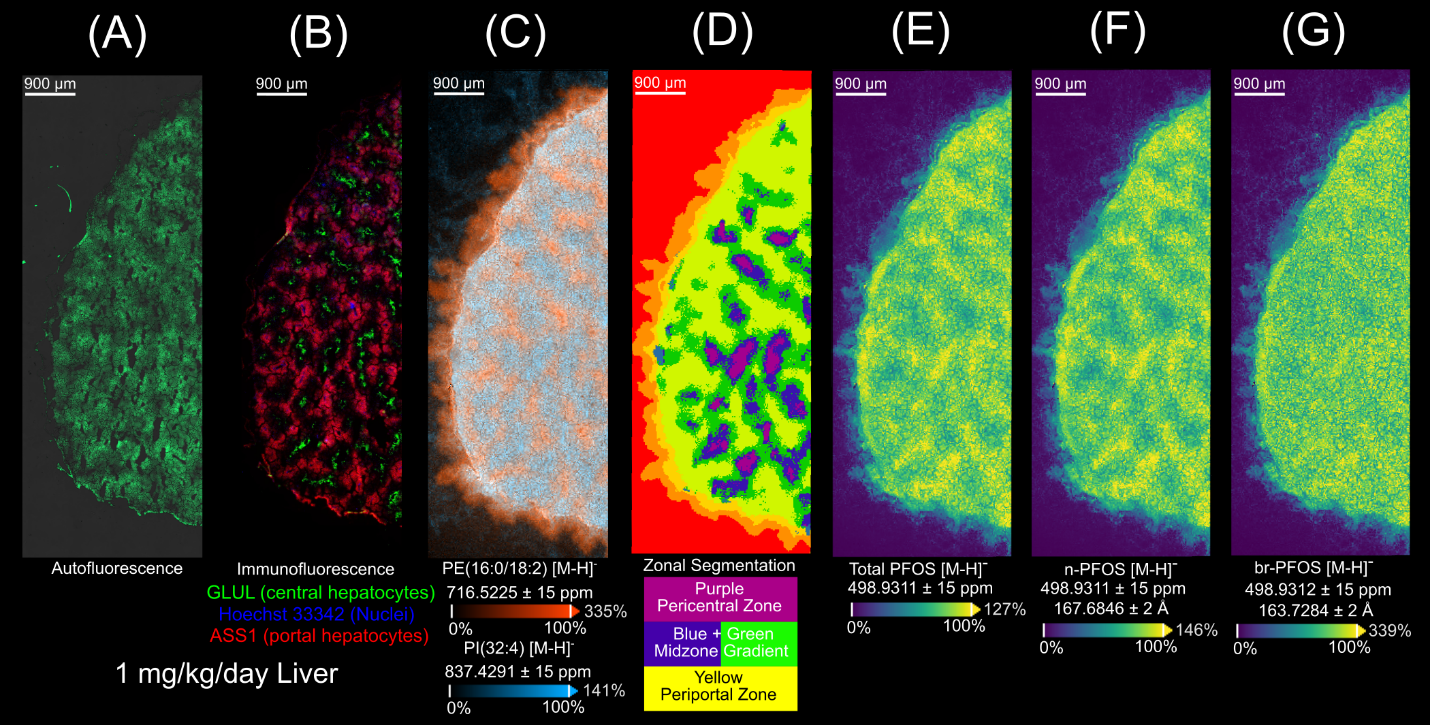

**Figure S7:** **Multi-modal imaging results on the same 1 mg/kg/day PFOS-exposed mouse liver tissue section.** (A) Pre-MALDI brightfield and GFP autofluorescence. (B) Post-MALDI immunofluorescence imaging of zonal hepatocyte markers. Livers were stained for Hoechst 33342 (Nuclei; blue), ASS1 (portal hepatocytes; red), and GLUL (central hepatocytes; green). (C) Periportal and pericentral distribution of PI(32:4) and PE(16:0/18:1), respectively, putatively annotated via accurate mass matching, and PE(16:0/18:1) confirmed with on-tissue MS/MS. (D) Zonal segmentation based on lipidomic profiles of imaged tissue regions visualizing pericentral zone (purple), midzone gradient (blue and yellow), and periportal zone (green). (E-G) MALDI-MSI ion images for total, linear and branched PFOS. MALDI-MSI parameters: matrix-DAN; polarity-negative; lateral spatial resolution-20 μm; other parameters-see Experimental.

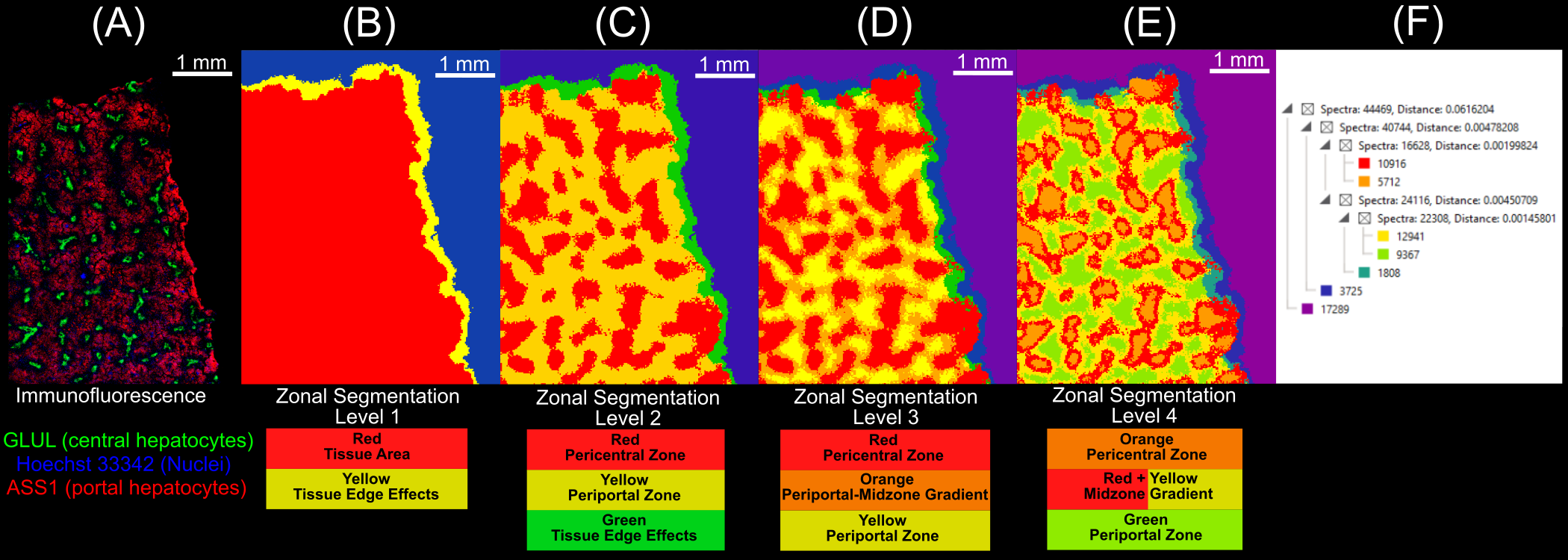
**Figure S8. Example segmentation analysis of a vehicle tissue towards accurate pericentral and periportal zonation.** (A) Immunofluorescence image of a vehicle control liver. Liver was stained for Hoechst 33342 (Nuclei; blue), ASS1 (portal hepatocytes; red), and GLUL (central hepatocytes; green). (B) First level segmentation defining tissue area (red) and tissue edge effects (yellow). (C) Second level segmentation further interrogating tissue are showing broad pericentral zones (red), periportal (yellow) and edge effects (green). (D) Third level segmentation interrogating periportal zone reveals periportal-midzone gradient (orange), defined periportal zone (yellow), and a broad pericentral zone (red). (E) Fourth level segmentation interrogating pericentral zone definition illustrating defined pericentral zones (orange), complete midzone gradient (red and yellow), and defined periportal zones (green). (F) The segmentation tree required to produce the segmented regions for accurate zonal areas.

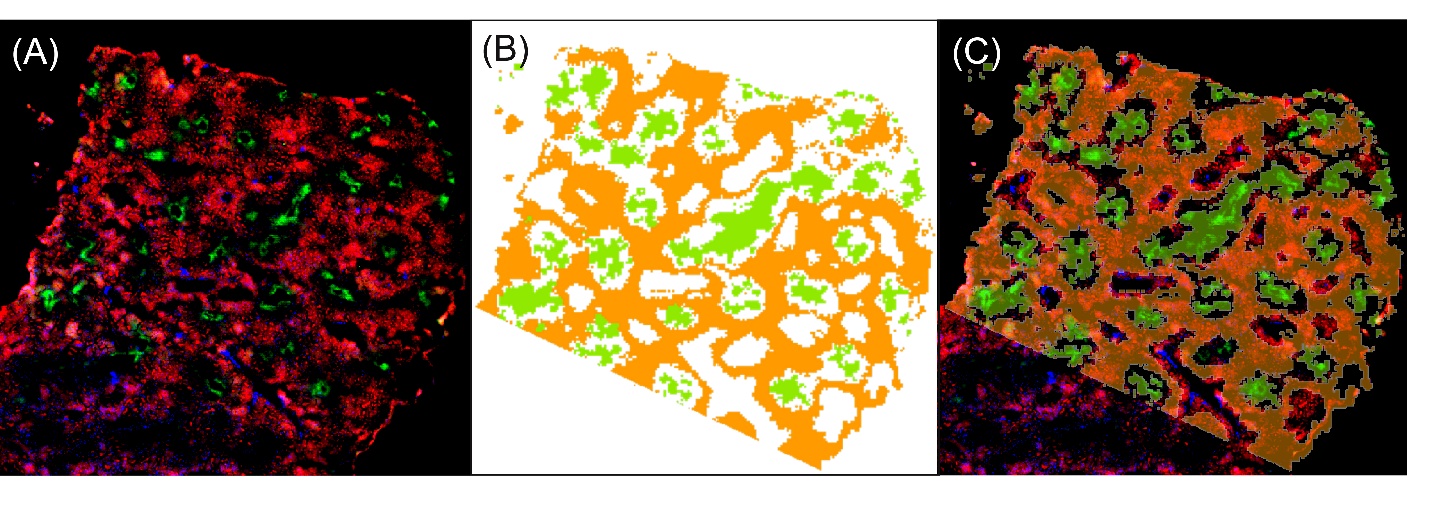

**Figure S9**. Example of validating segmented regions against immunofluorescence. (A) Post-MALDI immunofluorescence imaging of zonal hepatocyte markers. Livers were stained for Hoechst 33342 (Nuclei; blue), ASS1 (portal hepatocytes; red), and GLUL (central hepatocytes; green). (B) Extracted segmented pericentral (green) and periportal (orange) regions from MSI data. (C) Overlay of immunofluorescence and segmented regions, which shows co-localization of fluorescence markers and segmented regions.

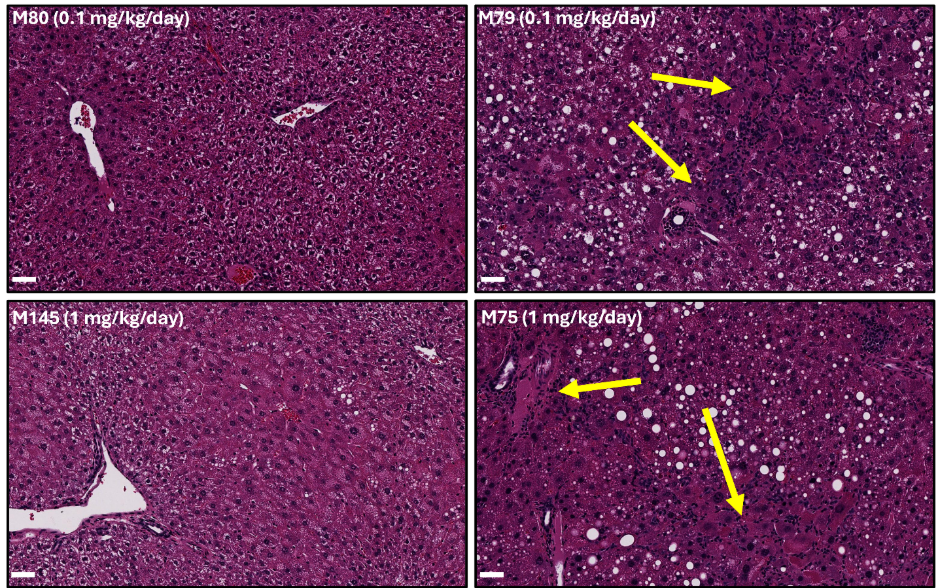

**Figure S10**. Representative photomicrographs of livers of PFOS treated mice showing expected pathological features in each treatment group (left), and abnormal pathologies compared to treatment group matched samples (right). Yellow arrows indicate regions of extensive cellular necrosis, hepatocyte hyperplasia, and inflammation that isn’t observed in other samples from the same treatment group. Scale bar represents 50 µm.

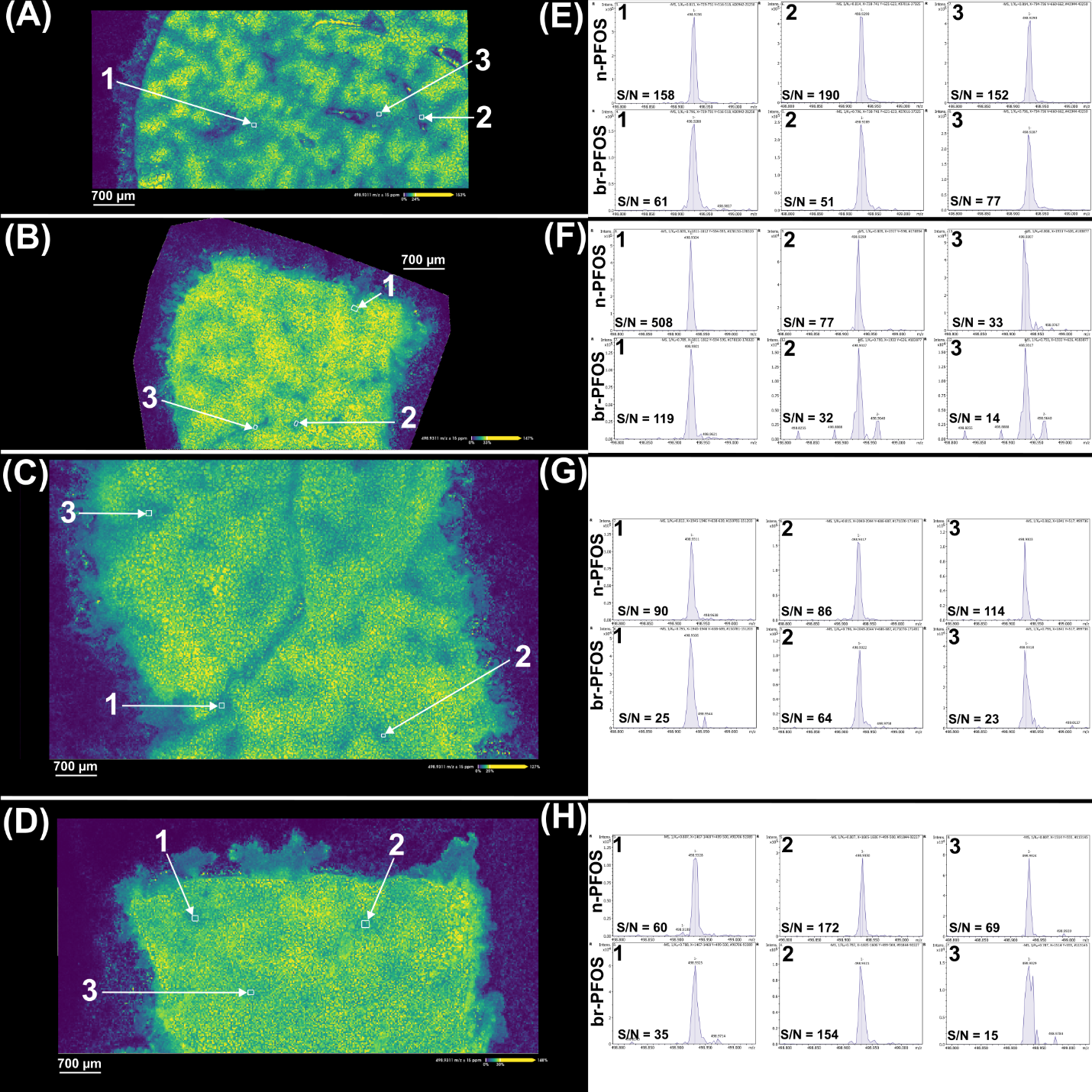

**Figure S11.** **Signal-to-noise ratio of PFOS isomers across low intensity liver regions.** (A-D) Ion images of 0.1 mg/kg/day treatment livers showing the total PFOS [M-H]^-^ distribution. Randomly selected areas of low PFOS [M-H]^-^ intensity regions within segmented periportal zones from imaged 0.1 mg/kg/day treatment PFOS livers were chosen for S/N calculations and are indicated by white arrows and corresponding numbers of 1, 2, or 3. (E-H) Spectra of n-PFOS and br-PFOS [M-H]^-^ isomers and their calculated S/N from areas 1, 2, or 3 as denoted in the spectra. Ion intensity scaling adjusted to better show PFOS [M-H]^-^ intensity distribution across low-dose tissues.

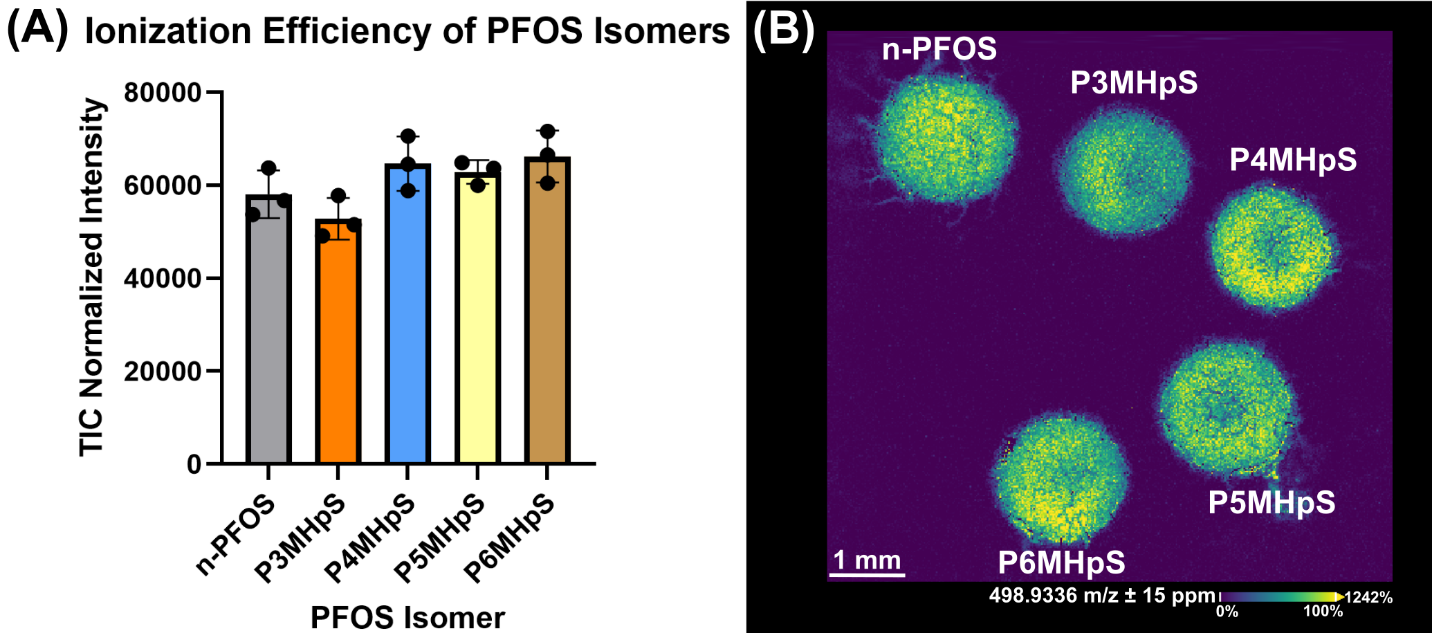

**Figure S12. On-tissue ionization efficiency of PFOS isomers by MALDI-TIMS-MSI.** (A) TIC normalized intensity of PFOS isomers spotted onto liver sections. Stock standards of 1 μg/mL were diluted with LC-MS grade water to yield 0.5 μg/mL solutions in 1:1 H2O:MeOH. 0.5 μL of each standard was spotted onto liver sections in technical replicates of n=3 for imaging. (B) Ion image of on-tissue spotted PFOS isomers to visualize similar intensity scaling.

**Figure S13**. Spatial analysis of PFOS zonal distributions for low-dose groups (0.1 mg/kg/day) using conditional autoregressive (CAR) modeling. Box plots show pixel-level distributions of the log2-transformed linear:branched PFOS intensity ratio in periportal and pericentral regions for individual liver sections. CAR models were fitted separately to each biological replicate to estimate the zonal effect while accounting for spatial autocorrelation among neighboring pixels.

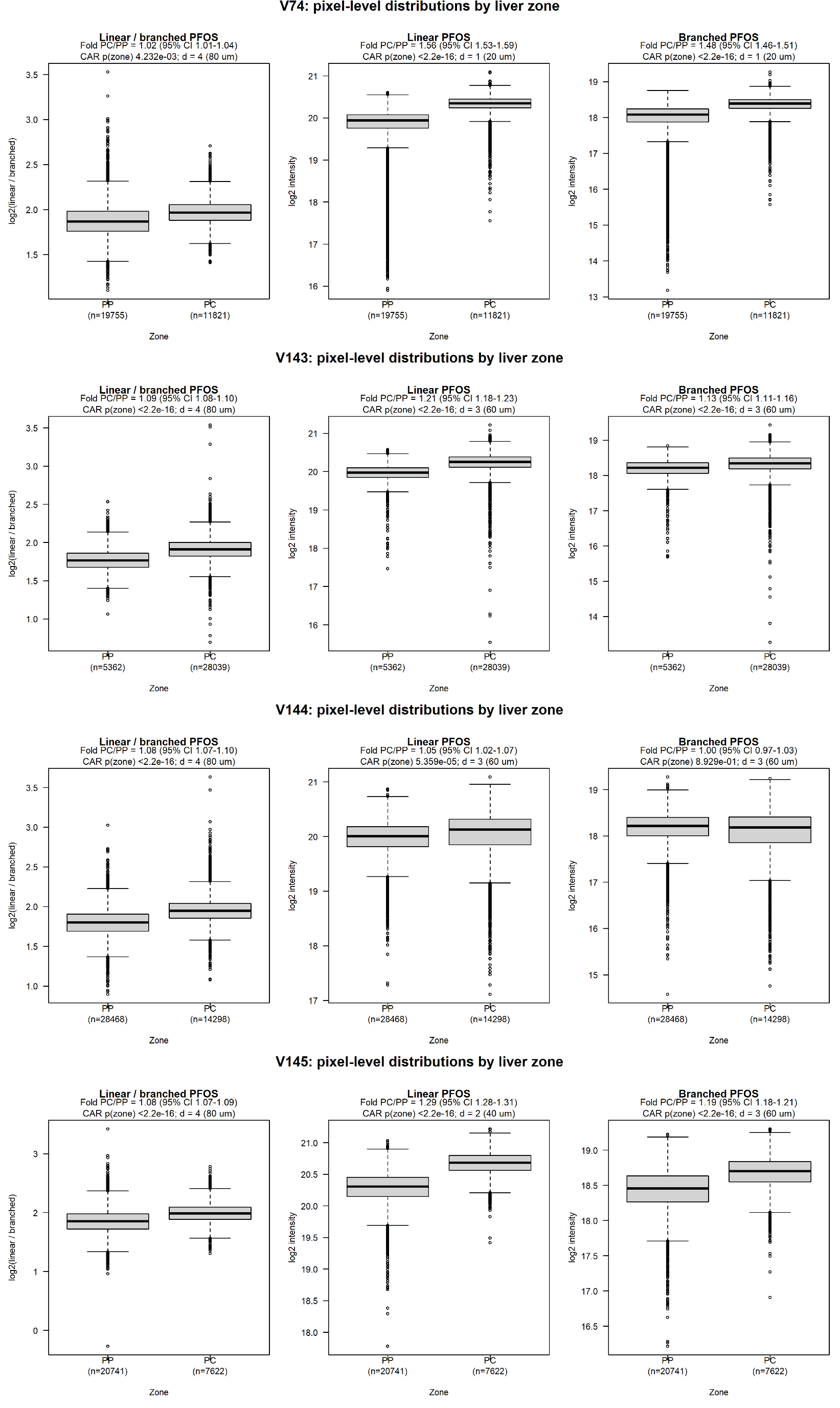

**Figure S14**. Spatial analysis of PFOS zonal distributions for high-dose groups (1 mg/kg/day) using conditional autoregressive (CAR) modeling. Box plots show pixel-level distributions of the log2-transformed linear:branched PFOS intensity ratio in periportal and pericentral regions for individual liver sections. CAR models were fitted separately to each biological replicate to estimate the zonal effect while accounting for spatial autocorrelation among neighboring pixels.

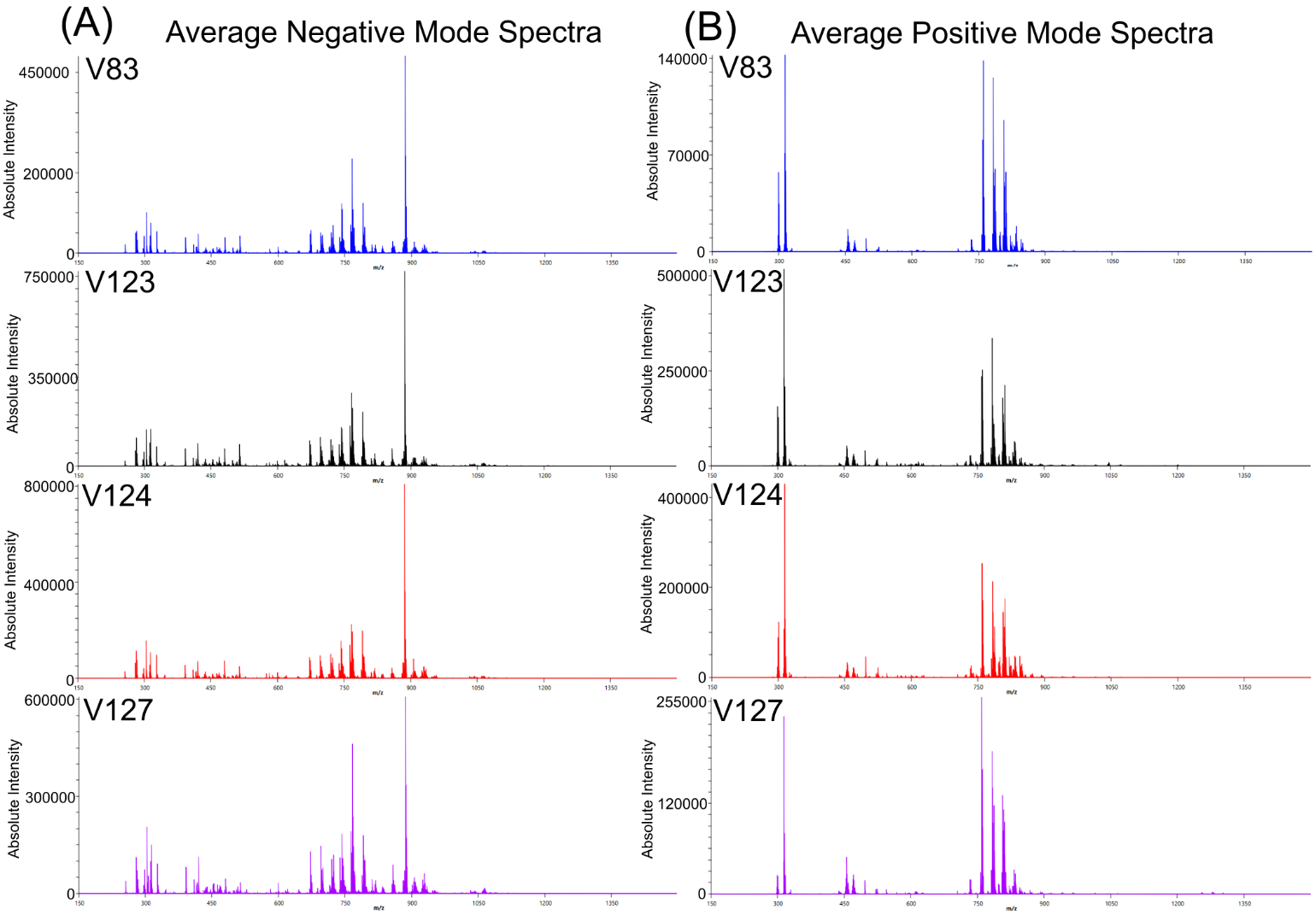

**Figure S15.** Average spectra of vehicle control tissues from negative mode and positive mode imaging. (A) Negative mode average spectra from imaged tissue areas. (B) Positive mode average spectra from imaged tissue areas.

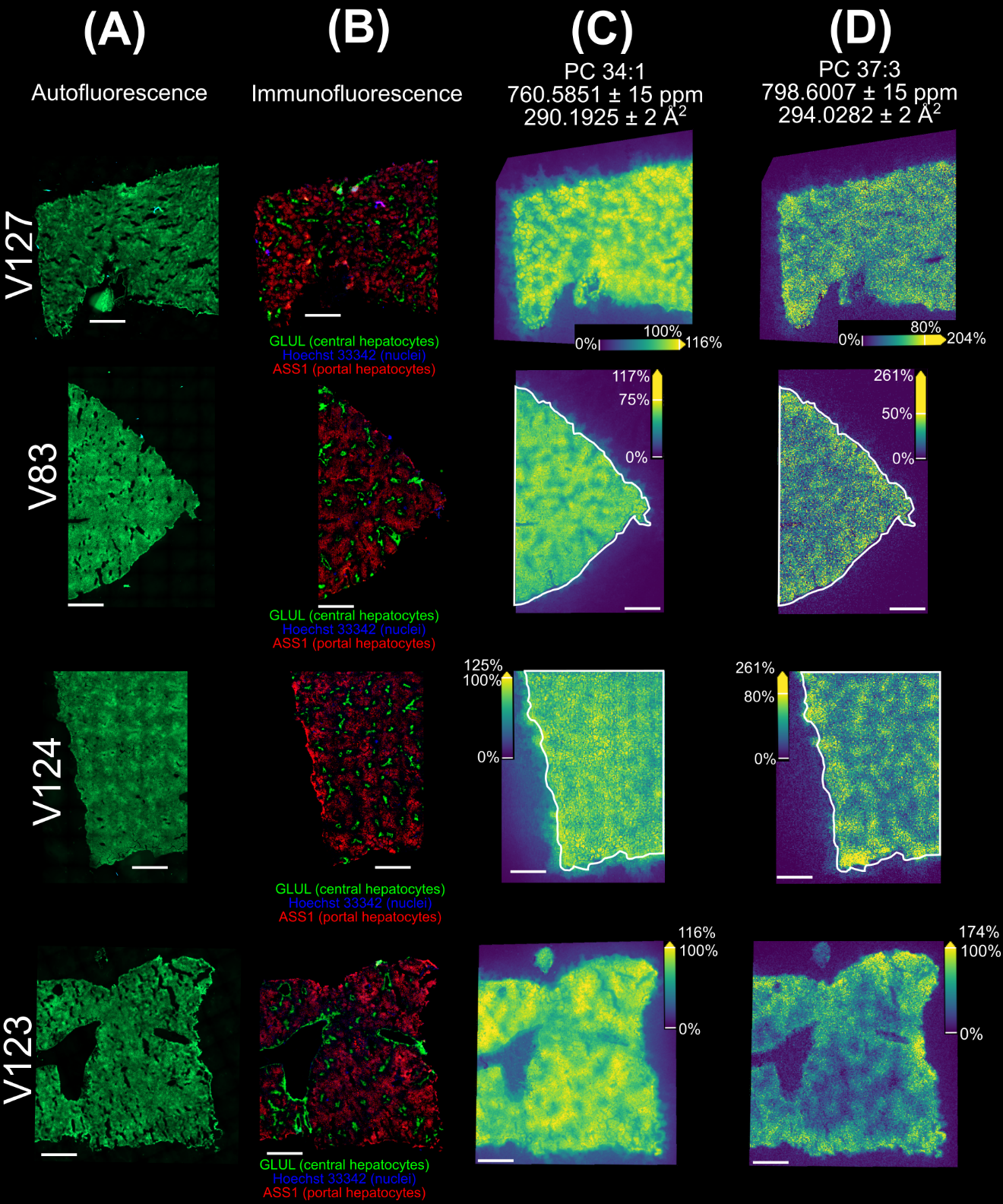

**Figure S16.** Positive mode lipid imaging visualizing a biomarkers of periportal and pericentral zones in vehicle control livers. (A) Pre-MALDI brightfield and GFP autofluorescence. (B) Post-MALDI immunofluorescence imaging of zonal hepatocyte markers. Livers were stained for Hoechst 33342 (Nuclei; blue), ASS1 (portal hepatocytes; red), and GLUL (central hepatocytes; green). (C-D) Periportal and pericentral distribution PC 34:1 and PC 37:3 respectively, putatively annotated via accurate mass matching and confirmed against existing literature.

**Table S3**. MS/MS annotations from iprm-PASEF analysis. More information can be found in Supplemental Materials.

| **Tentative Lipid + Precursor m/z** | **MS/MS Identities** |
| --- | --- |
| LPI 18:0 m z 599.3202 | LPI 18:0 |
| LPI 20:4 m z 619.2889 | LPI 20:4 |
| PA 32:0 m z 647.4647 | PA 16:0/16:0 |
| PA 34:2 m z 671.4657 | PA 16:0/18:2 |
| PA 34:1 m z 673.4814 | PA 16:0/18:1 |
| PA 36:4 m z 695.4657 | PA 16:0/20:4, PA 18:2/18:2 |
| PA 36:3 m z 697. 4814 | PA 16:0/20:3, PA 18:1/18:2 |
| PA 36:2 m z 699.4970 | PA 18:1/18:1 |
| PE 34:2 m z 714.5100 | PE 16:1/18:1, PE 16:0/18:2 |
| PE 34:1 m z 716.5236 | PE 16:0/18:1, PE 16:1/18:0 |
| PA 38:6 m z 719.4657 | PA 18:2/20:4 |
| PA 38:5 m z 721.4814 | PA 16:0/22:5, PA 18:2/20:3, PA 18:1/20:4 |
| PA 38:4 m z 723.4970 | PA 18:1/20:3, PA 18:0/20:4, PA 16:0/22:4 |
| PA 38:3 m z 725.5127 | PA 18:0/20:3, PA 18:1/20:2, PA 18:2/20:1 |
| PE 35:2 m z 728.5236 | PE 17:1/18:1, PE 16:1/19:1 |
| PE 36:4 m z 738.5079 | PA 16:0/20:4 |
| PE 36:3 m z 740.5236 | PE 16:0/20:3 |
| PE 36:2 m z 742.5392 | PE 18:0/18:2. PE 16:0/20:2 |
| PE 36:1 m z 744.5549 | PE 18:0/18:1 |
| PA 40:6 m z 747.4970 | PE 18:1/22:5, PE 18:0/22:6 |
| PA 40:5 m z 749.5117 | PA 18:0/22:5, PA 18:1/22:4, PA 20:1/20:4 |
| PE 38:6 m z 762.5079 | PE 18:1/20:5, PE 16:0/22:6 |
| PE 38:5 m z 764.5236 | PE 18:1/20:4, PE 16:0/22:5 |
| PE 38:4 m z 766.5392 | PE 16:0/22:4, PE 18:0/20:4 |
| PE 38:3 m z 768.5549 | PE 18:2/20:1, PE 18:0/20:3 |
| PE 38:2 m z 770.5705 | PE 18:0/20:2, PE 18:1/20:1 |
| PE 40:6 m z 790.5392 | PE 18:0/22:6, PE 18:1/22:5 |
| PE 40:5 m z 792.5549 | PE 18:0/22:5 |
| PE 40:4 m z 794. 5705 | PE 20:0/20:4 |
| PE 40:3 m z 796.5862 | PE 20:0/20:3, PE 18:0/22:3 |
| PS 38:4 m z 810.5291 | PS 18:0/20:4 |
| PE 42:8 m z 814.5392 | PE 20:2/22:6 |
| PE 42:7 m z 816.5549 | PE 20:1/22:6 |
| PE 42:6 m z 818.5705 | PE 20:0/22:6 |
| PE 42:5 m z 820.586 | PA 20:0/22:5, PA 20:1/22:4 |
| PI 34:2 m z 833.5186 | PI 16:0/18:2 |
| PS 40:6 m z 834.5291 | PS 18:0/22:6 |
| PS 40:5 m z 836.5447 | PS 18:0/22:5 |
| PI 36:3 m z 859. 5447 | PI 18:1/18:2 |
| PI 36:2 m z 861.5499 | PI 18:1/18:1 |
| PI 38:5 m z 883.5342 | PI 18:1/20:4 |

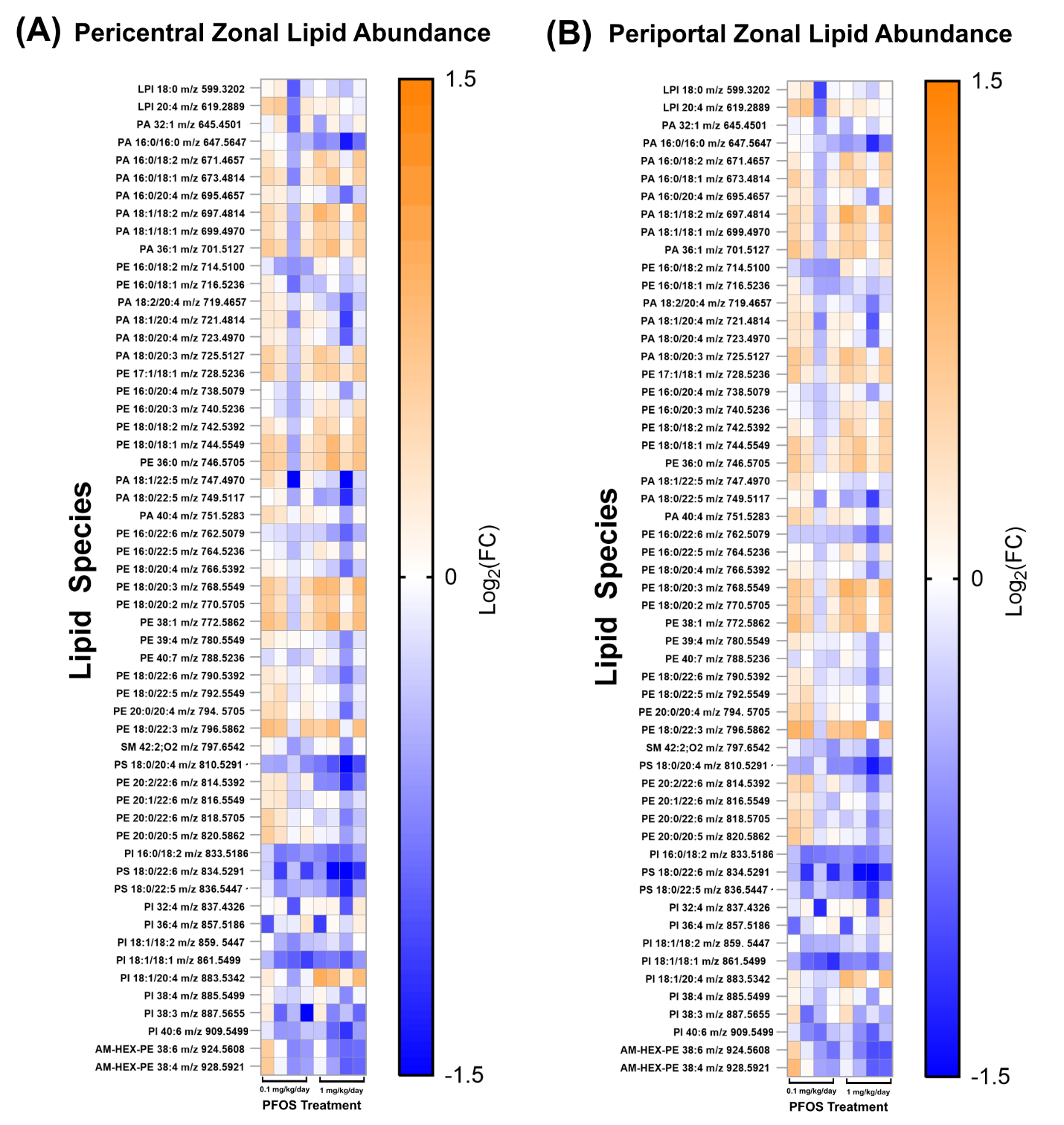

**Figure S17: Heatmaps illustrating individual replicate response and lipid abundance fold change to PFOS treatment.** (A) Abundance fold change of lipid species in pericentral zones of mouse livers. (B) Abundance fold change of lipid species in periportal zones of mouse livers. Columns represent individual replicates from both 0.1 mg/kg/day and 1 mg/kg/day treatment groups. MALDI-MSI parameters: matrix-DAN; polarity-negative; lateral spatial resolution-20 μm; other parameters-see Experimental methods.

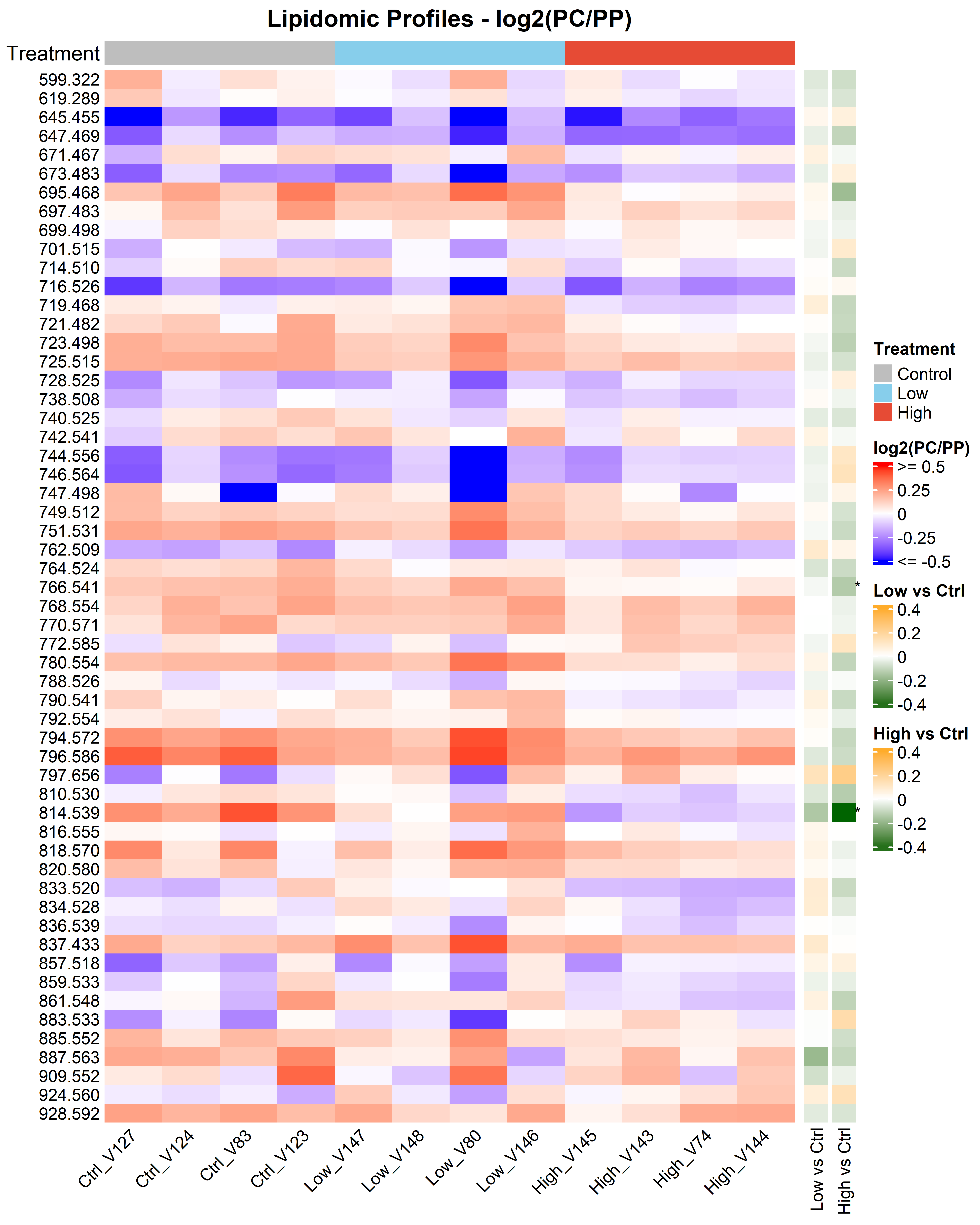

**Figure S18**. Heatmap of zonal ratios for lipids and treatment groups. For each lipid feature, zonal distribution was represented as the log2-transformed ratio of PC:PP intensities. Positive values (red scale) indicate pericentral preference. Negative values (blue scale) indicate periportal preference. Zero indicates no preference. Treatment groups are indicated by different colors. For each lipid, effects of treatment on log2(PC:PP) ratios were evaluated using Dunnett’s multiple comparison to compare the low- and high-dose groups separately to the control and Benjamini-Hochberg FDR correction for lipids. Fold changes (low vs ctrl, high vs ctrl) were shown as color bars near the clustering map, and statistical significance was defined as FDR-adjusted q < 0.05 and shown near the fold change bars.

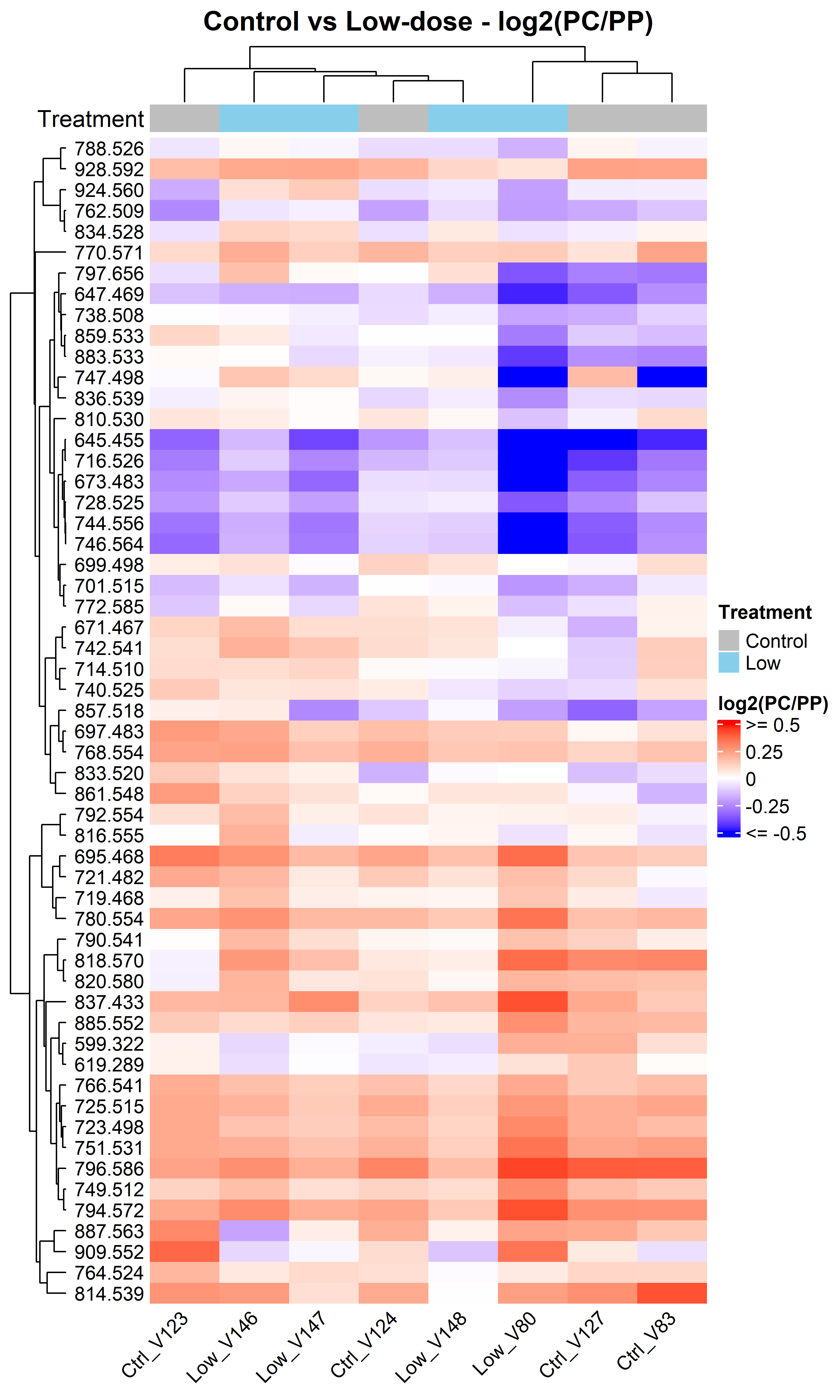

**Figure S19**. Hierarchical clustering with control and low-dose group only.

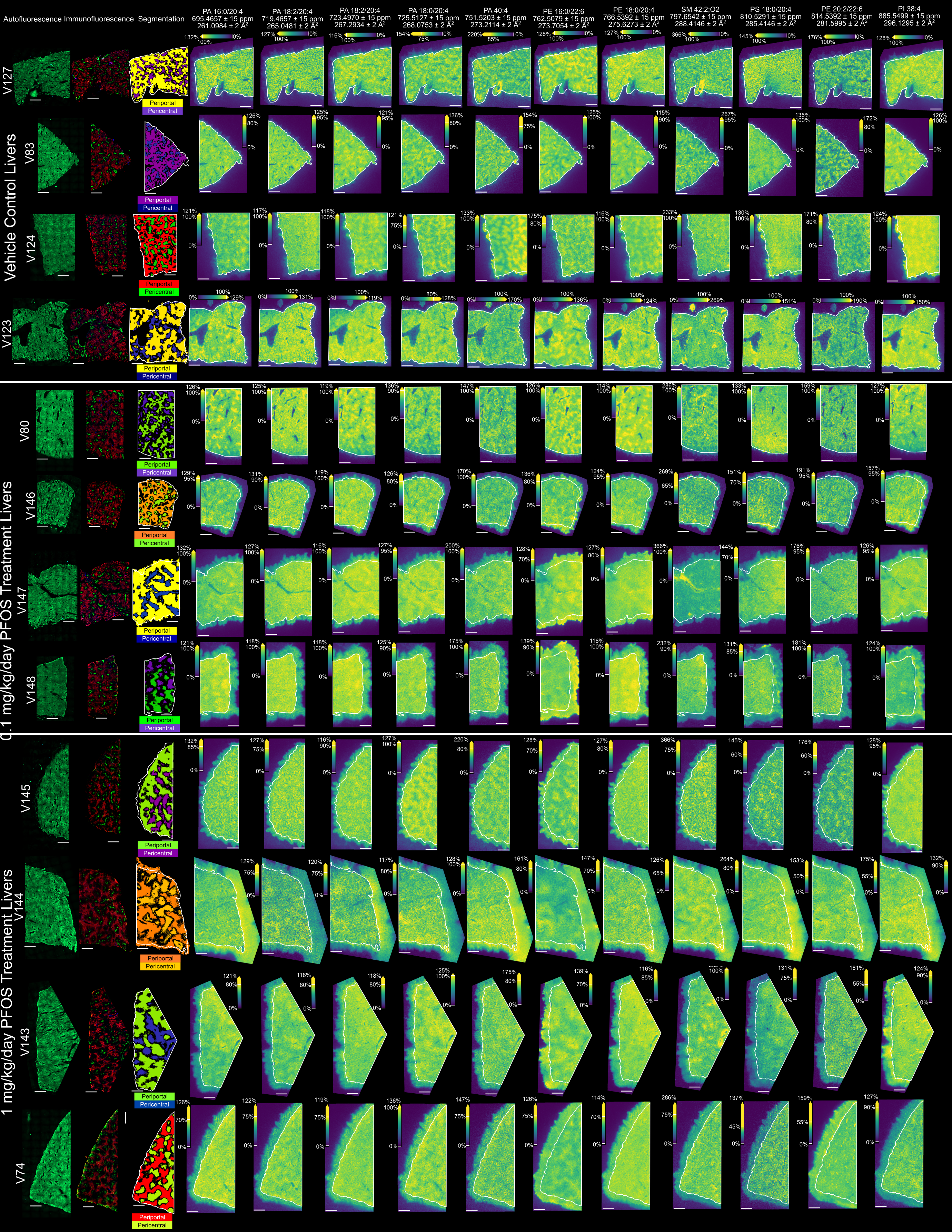

**Figure S20:** Comprehensive lipid ion images showing 11 selected lipids. Ion image intensities are adjusted per image to create similar color intensity scaling for ease of visualization of zonal localization across treatment conditions. Outlines of tissues in segmentation maps and ion images are indicated by white border. White scale bars indicate 900 μm.
